# Physicochemical changes in plasma membrane mirror nanoparticle-mediated cytotoxicity

**DOI:** 10.1101/2019.12.29.890236

**Authors:** Vishesh Sood, Dhirendra S. Katti

## Abstract

The aim of this chapter was to understand the influence of nanoparticle challenge on physicochemical characteristics of the cells and to correlate these changes with cytotoxic response of nanoparticles. A nanoparticle surface charge and a concentration-dependent cytotoxic response were observed in the breast cancer cell lines MDA MB 231 and SKBR3. The cationic gold nanoparticles were more cytotoxic to cells as compared to anionic gold nanoparticles. It was also observed that cationic nanoparticles compromised the integrity of the plasma membrane at higher concentrations. Cationic nanoparticle challenge also caused changes in physicochemical characteristics of plasma membrane. Higher concentration of cationic nanoparticles caused an irreversible change in the surface charge density of cells. However, anionic gold nanoparticles did not show any such effect. It was observed that the ROS-mediated oxidative stress was the mechanism of cationic gold nanoparticle-mediated cytotoxic effect. Mitochondrial depolarization was observed in both anionic and cationic nanoparticle challenge. Therefore, the role of mitochondrial ROS in nanoparticle-mediated cytotoxicity is questionable. Finally, a generalized model involving modulation of intracellular Ca^2+^ can potentially provide an explanation for the observed pluralistic response of the cells towards nanoparticle challenge.

## Introduction

Nanoparticle-mediated cellular responses are instrumental in defining the biological activity of nanoparticles and, therefore, are important for the safe and efficacious use of nanoparticles in biomedical applications. Nanoparticles, owing to their small size, have a very high surface energy and, therefore, can interact with the biomacromolecules and cells present in their vicinity. These interactions mediate the biological response elicited by nanoparticle challenge. Physicochemical characteristics of nanoparticles such as size, shape, surface charge, and bulk material affect their biological activity^1^. Nanoparticles, however, do not interact with the cells directly. Biomacromolecules present in the medium interact with the nanoparticles, and this interaction forms a biomolecular corona around the nanoparticles^2,3^. The biomolecular corona is primarily formed by proteins although other biomacromolecules such as lipids^4^ and sugars^5^ are also present. The biomolecular corona provides a unique biological identity to the nanoparticles and mediates the interactions between nanoparticles and cells^6^. Nanoparticles are known to elicit a cell type and bulk material specific cellular response. For example, gold nanoparticles (AuNP) can cause a cytotoxic response based on cell surface charge^7^, on the other hand, they can also enhance the osteogenic differentiation of mesenchymal stem cells^8^. There is no model available in literature that can explain the plurality of the nanoparticle-mediated cellular response.

To understand the fundamental nature of nanoparticle-mediated cellular responses, it is imperative to understand the nature of nanoparticle-cell interface. *In vitro* nanoparticle-cell interface is formed between the nanoparticles suspended in cell culture media and cells seeded on a polystyrene substrate. However, the nanoparticles perceived by cells are not pristine or as synthesized. Physicochemical characteristics of nanoparticle change on exposure to cell culture media^9^. The high ionic strength of medium causes agglomeration of nanoparticle resulting in an increase in the size of nanoparticles^10^. Simultaneously, a biomolecular corona is formed around the nanoparticles which help them to stay in a dispersed phase. These two opposing forces result in a new state of nanoparticle with altered physicochemical characteristics. Agglomeration of nanoparticles influences the biological behavior of cells. For example, agglomerated AuNP cause enhanced cell death when compared to monodispersed AuNP^10^. Moreover, agglomerated AuNP also cause activation of autophagy in the cells^11^. Therefore, the physicochemical characteristics of nanoparticles once dispersed in cell culture media have a direct correlation with the biological response.

Cells, which form the other part of nanoparticle-cell interface, do not act as a passive bystander. Cells actively internalize nanoparticles as a consequence of nanoparticle-cell interaction^12^. For nanoparticle internalization, nanoparticle-cell interaction should be able to compensate for the instability of the membrane caused by bending of the plasma membrane, and exposure of hydrophobic core of phospholipid bilayer of the plasma membrane caused as a result of membrane wrapping around nanoparticles^12,13^. In an *in vitro* setup, cells are adhered to a rigid substrate and therefore have an endocytic hot-spots present on their apical surface. Nanoparticles tend to interact with the plasma membrane, and then nanoparticles are actively taken to the endocytic hot-spot while still bound to the plasma membrane. Therefore, only those nanoparticles are able to reach apical surfaces which have a strong affinity to the cell surface^14^. For uptake of nanoparticles by cells, the overall interaction energy between nanoparticles and plasma membrane biomolecules such as lipids, glycoproteins, and cell surface receptors, should be able to compensate the energy loss due to plasma membrane destabilization. Therefore, nanoparticle-plasma membrane interactions form the basis of the nanoparticle-cell interface^12^. Nanoparticle-cell interaction results in the physicochemical changes in the structure of plasma membrane. For example, cationic AuNP on interaction with cells causes a significant change in the plasma membrane potential^7^. On the contrary, anionic and neutral AuNP do not cause any change. Moreover, nanoparticle incubation with cells also causes a change in the zeta potential of cells which indicates that the overall composition of the plasma membrane is changing as a result of nanoparticle internalization^15^.

The plasma membrane is a dynamic organelle of the cell and is actively involved in sensing environmental stress, and it also plays a role in cellular signaling. Change in the plasma membrane fluidity acts as a signal for temperature fluctuations in the environment. The increase in environmental temperature causes a corresponding increase in the lateral fluidity of the plasma membrane. These changes are known to cause activation of classical heat shock response^16-18^. Moreover, hypopolarization of the plasma membrane results in the activation of MAP-kinase leading to cellular proliferation^19^. Collectively, these results point towards activation of different cellular pathways as a result of changes in the physicochemical properties of the plasma membrane. Based on this observation, ***we hypothesized that changes in physicochemical properties of the plasma membrane as a consequence of nanoparticle challenge could explain the plurality of the responses observed during the nanoparticle-cell interaction.*** To test this hypothesis, breast cancer cell lines (MDA MB 231 and SKBR3) were challenged with anionic and cationic AuNP to evaluate the cytotoxic response of cells. Further, in this study the physicochemical changes in the plasma membrane structure were assessed after nanoparticle challenge to study the link between nanoparticle-cell interaction and cytotoxic response. Taken together, results presented in this chapter points towards a generalized mechanism of cytotoxic response of nanoparticles, wherein, nanoparticles alter the physicochemical structure of plasma membrane upon their internalization by cells and these changes are perceived by the cells as an environmental stress. Finally, a generalized nanocytotoxicity model was proposed which is based on physicochemical changes in the plasma membrane caused by the nanoparticle-cell interaction. Once validated, this model can help in explaining the plurality in the nanoparticle-mediated responses of different cells.

## Materials and methods

### Synthesis of gold nanoparticles (AuNP)

#### Synthesis and characterization of anionic AuNP

Citrate reduction method^13^ was modified for the synthesis of quasi-spherical citrate-capped AuNP. Briefly, 49.55 mL of trisodium citrate solution (5.4 mM) was heated at 100 °C for 10 minutes under reflux conditions. The solution was stirred using a magnetic stirrer (RCT Basic, IKA^®^, India). To this 150 µL of HAuCl_4_ solution (100 mM) was added and allowed to react for 10 minutes. Following this, the temperature of oil bath was reduced to 90 °C followed by the addition of a second aliquot of 150 µL of HAuCl_4_ solution (100 mM) to the reaction. The reaction was then allowed to proceed for 20 minutes. Finally, a third aliquot of 150 µL of HAuCl_4_ solution (100 mM) was added to the reaction. The reaction was further allowed to proceed for another 20 minutes. After completion of the reaction, the oil bath was removed and AuNP suspension cooled for 30 minutes under mild stirring at room temperature. The final suspension of citrate-capped AuNP was transferred to a sterile polypropylene tube and stored at 4 °C until further use.

#### Synthesis and characterization of cationic AuNP

Cationic or amine-capped AuNP were prepared using a modified sodium borohydride reduction method. Briefly, 450 µL of HAuCl_4_ solution (100 mM) and 225 µL of aminoethanethiol (500 mM) was added to 49.075 mL of type I water. The reaction mixture was mildly stirred at room temperature for 30 minutes followed by the addition of 250 µL of ice-cold NaBH_4_ solution (100 µM). The reaction was allowed to continue for 15 min at room temperature. Upon the completion of the reaction, the reaction mixture was transferred to a polypropylene tube and kept undisturbed at room temperature for 24 hours. The synthesized AuNP suspension was stored in a sterile polypropylene tube at 4 °C until further use.

### Characterization of AuNP

#### Morphology

The morphology of as-prepared AuNP was studied using a transmission electron microscope. For TEM samples, the nanoparticle suspension was dried on a glass slide for 2 hours at room temperature. The precipitate from the edge of the drop was collected on a Formvar-coated 300 mesh copper grid (Tedpella Inc., USA). The samples were then analyzed using a Tecnai G2 12 Twin TEM (FEI, USA) operating at 120 kV.

#### Size

The hydrodynamic diameter of AuNP was estimated by photon correlation spectroscopy using a Zetasizer ZS90 (Malvern Instruments Ltd., UK) fitted with a laser diode of 633 nm wavelength. Briefly, 1 mL sample of as prepared AuNP suspension was added to a disposable cuvette, and the sample was equilibrated at 25 °C for 180 seconds. The time-dependent decay in the intensity of light was measured and converted into a correlogram that was used to estimate the hydrodynamic diameter of AuNP using the general mode of analysis in the Zetasizer software.

#### Zeta potential

The zeta potential of as-prepared AuNP was estimated by Laser Doppler Velocimetry (LDV) using Zetasizer ZS90. LDV measures electrokinetic behavior of nanoparticles under an externally applied electric field.

#### Nanoparticle suspension preparation

For all studies, AuNP dilutions were prepared by adding 450 µL of as prepared AuNP suspension into 550 µL of 2x cell culture media (1:1 mixture of Ham’s F12 nutrient mixture and DMEM) supplemented with 10% FBS. To control agglomeration of nanoparticles, the suspension was thoroughly mixed by repeated aspiration and dispensation using a pipette.

#### Cell culture protocol

MDA MB 231 and SKBR3 were the two breast cancer cell lines used for this study. Both cell lines were derived from the pleural effusion from the metastatic site of adenocarcinoma of the mammary gland of Caucasian females. The culture media used for growing cells was 1:1 mixture of Ham’s F12 nutrient mixture and DMEM supplemented with 10% FBS. All experiments were performed within five passages of the cells.

#### Height of culture media

The height of media in a multiwell plate (*h*) was calculated as the ratio of the difference in the optical density at 900 and 977 nm of the media to that of blank phases measured in a 1 cm reference cell in Take 3^®^ plates (equation 1).

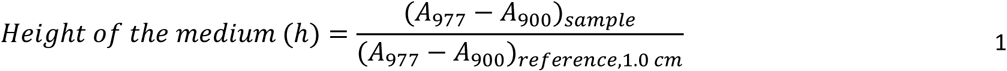

#### Cell viability assay

Resazurin assay was used to study the effect of anionic and cationic AuNP challenge on the viability of MDA MB 231 and SKBR3 cells^20^. Briefly, cells were seeded at a density of 15,625 cells cm^−2^ in a 96 well plate. After 24 hours, cells were gently washed with phosphate buffered saline without Ca^2+^ and Mg^2+^ ions (PBS^−^). Cells were then challenged with different dilutions of anionic and cationic AuNP prepared from the nanoparticle suspension as per section 0. Cells were challenged with nanoparticles for 24 and 48 hours. At the end of the challenge, media was removed, and cells were gently washed with phosphate buffered saline with Ca^2+^ and Mg^2+^ ions (PBS^+^). The cells were then incubated with resazurin assay reagent (0.02 mg mL^−1^ concentration of resazurin in complete cell culture media) for 5 hours, and the fluorescence was measured at an excitation wavelength of 540 nm and an emission wavelength of 600 nm using Synergy H4 multimode reader (Biotek, USA). The results were reported as percentage viability with respect to the untreated control.

#### Plasma membrane integrity assay

Plasma membrane integrity of cells after AuNP challenge was measured by fluorescein diacetate (FDA) and propidium iodide (PI) dual staining^21^. Briefly, MDA MB 231 cells were seeded at a density of 15,625 cells cm^−2^ in a 48 well plate followed by 24 hours of incubation. Cells were then challenged with 20 µg mL^−1^ and 80 µg mL^−1^ concentration of anionic and cationic AuNP for 3 and 12 hours. After incubation media was discarded and cells were gently washed with PBS^+^. Cells were then incubated with freshly prepared FDA/PI staining solution (5 mL complete media without FBS, 8 µL of the 5 mg mL^−1^ stock solution of fluorescence diacetate prepared in acetone, and 50 µL of the 2 mg mL^−1^ stock solution of propidium iodide prepared in PBS). The cells were incubated for 5 minutes at room temperature, and images were captured using a fluorescent microscope (Nikon Eclipse TE2000-U).

#### Cell surface charge measurement

The cell surface charge density was estimated by measuring the electrophoretic mobility of the cells under an applied electric field by Laser Doppler Velocimetry using Zetasizer ZS90 (Malvern Instruments Ltd., USA). The zeta potential of cells was calculated from the electrophoretic mobility data. The suspending medium used for zeta potential measurement was NaCl solution. The ionic strength of the medium was optimized to avoid interference of ions with the zeta potential measurement. For this, the zeta potential of MDA MB 231 cells was measured in media of varying ionic strength while maintaining the osmolality at 380 mOs using glucose. Finally, the influence of nanoparticle challenge on surface charge density of AuNP was studied. Briefly, cells were seeded at a density of 15,625 cells cm^−2^ in a 24 well plate and were exposed to different concentrations of anionic and cationic AuNP for 3 and 6 hours. After incubation, media was removed, and cells were gently washed using PBS^+^. Cells were detached from the substrate by using non-enzymatic cell dissociation reagent (SIGMA-Aldrich, USA) in order to avoid harsh trypsin-EDTA which alters the cell surface charge by lysing cell surface proteins.

#### Plasma membrane fluidity assay

Lateral diffusivity of the plasma membrane was determined using pyrene excimerization assay. Pyrene is a polycyclic organic compound which shows concentration-dependent excimerization and the rate of excimer formation is diffusion dependent and thus is a function of the viscosity of the surrounding media^22^. Briefly, after nanoparticle treatment of cells for desired time, cells were washed gently with PBS^+^. Cells were incubated with the pyrene reagent (4 µM pyrene solution in PBS^−^ with 0.08% w/v Pluronic^®^ F127) for 10 minutes in a CO_2_ incubator. The amount of pyrene monomers left was estimated by measuring the fluorescence intensity of pyrene monomer at an excitation wavelength of 340 nm and an emission wavelength of 380 nm using Synergy H4 multimode reader (Biotek, USA). The results were reported as a percentage change in the plasma membrane fluidity with respect to the untreated control.

#### Mitochondrial membrane potential assay

Mitochondrial membrane potential was measured using rhodamine 123 (R123) accumulation assay^23^. R123 is a voltage sensitive probe which accumulates inside the mitochondria. Briefly, MDA MB 231 cells were seeded at a density of 15,625 cells cm^−2^ in a 24 well plate. After 24 hours of incubation, cells were challenged with different concentrations of anionic and cationic AuNP for 3 hours. The media was removed, and cells were gently washed with PBS^+^. Cells were incubated with R123 reagent (1 µM R123 suspended in cell culture medium) for 15 minutes in a CO_2_ incubator followed by gentle washing with PBS^+^. The accumulation of R123 in the cells was measured using area scan mode of Synergy H4 multimode reader (Biotek, USA). The emission wavelength used was 510 nm and excitation wavelength used was 534 nm with 9 nm bandpass. All results were reported as a percentage change in the accumulation of R123 with respect to the untreated control.

#### Intracellular Reactive Oxygen Species assay

Intracellular ROS was measured using dichlorofluorescein diacetate (DCF-DA) assay^24^. Briefly, MDA MB 231 cells were seeded at a density of 15,625 cells cm^−2^ in a 24 well plate and incubated for 24 hours. The cells were challenged with different concentrations of anionic and cationic AuNP for 3 hours. After the AuNP challenge, media was removed, and cells were washed gently with PBS+. Cells were then incubated with DCF-DA reagent (100 µM DCF-DA in 1:1 mixture of Ham’s F12 and DMEM supplemented with 1% FBS) and incubated in a CO_2_ incubator. The fluorescence intensity was measured using Synergy H4 multimode reader (Biotek, USA) maintained at 37°C during measurement. The excitation wavelength used was 485 nm and the emission wavelength used was 530 nm with 9 nm bandpass. The results were analyzed using equation 2.

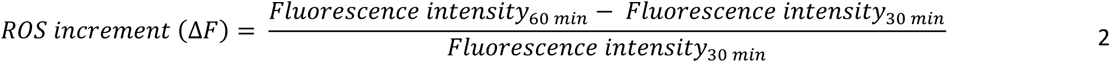

## Results and discussion

### Synthesis and characterization of gold nanoparticles (AuNP)

Reported methods for the synthesis of anionic and cationic AuNP were modified to synthesize concentrated suspension of nanoparticles as described in section 0. The morphology of synthesized AuNP was quasi-spherical as observed using transmission electron microscopy (**Figure 1 A** and B). Nanoparticles were polycrystalline in nature as indicated by the small area X-ray diffraction pattern (**Figure 1 A** and B, inset).

**Figure 1.**
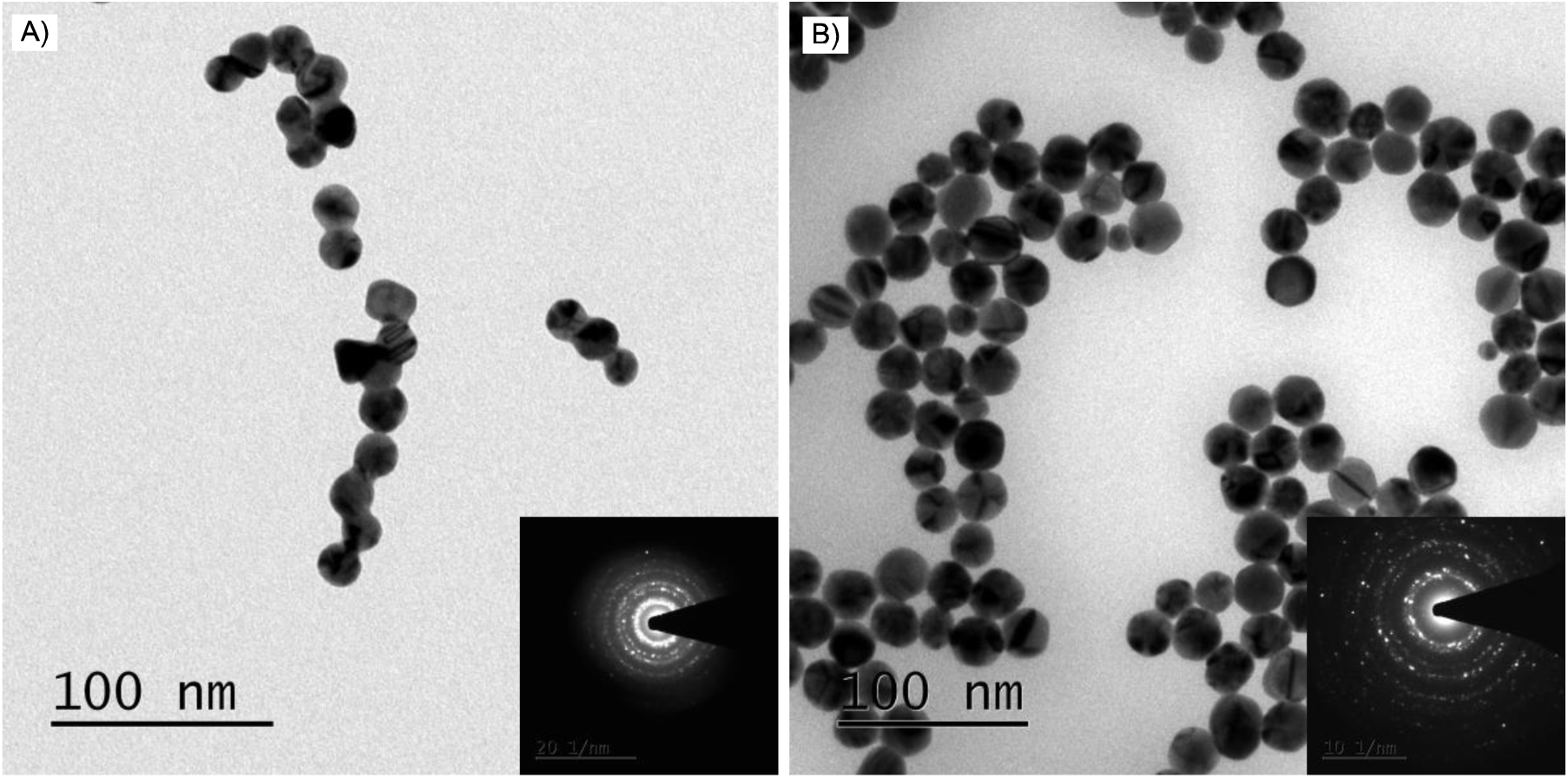
Transmission electron micrographs of nanoparticles. A) Anionic AuNP, and B) Cationic AuNP (scale bar – 100 nm). Inset figures show selected area diffraction patterns of anionic and cationic AuNP in the respective images.

The size of nanoparticles was measured using Photon correlation spectroscopy (PCS). The Z-average of hydrodynamic diameter of citrate-capped AuNP estimated by PCS had a value of 23.17 nm and a PDI of 0.046 (**Figure 2A**). The nanoparticle suspension was free from any large agglomerates as is evident from cumulative intensity curve (**Figure 2A**) and correlogram of photon decay (**Figure 2A A**, inset). The value of 95% undersize was 36.4 nm and that of 50% undersize was 23.9 nm. The Z-average of hydrodynamic diameter of AET-capped AuNP was 23.96 nm with a PDI of 0.144 as estimated by PCS (**Figure 2B**). For AET-capped nanoparticles, the value of 95% undersize was 44.4 nm and that of 50% undersize was 25 nm. These results indicate that the synthesized AuNP were monodisperse and were of comparable size. The surface charge density of synthesized nanoparticles was estimated by measuring their zeta potential using Laser Doppler Velocimetry (LDV). The zeta potential of citrate-capped nanoparticles was -32.40 ± 2.07 mV while that of AET-capped nanoparticles was 22.50 ± 2.57 mV. These results indicate that citrate-capped AuNP were anionic, whereas, AET-capped AuNP were cationic in nature. Moreover, the zeta potential distributions were monomodal (**Figure 3A**) indicating absence of differentially ionizing functional groups on the surface of nanoparticles. Also, the phase plots of LDV (**Figure 3B**) showed no significant shift of the baseline indicating that the sample was not charging during the measurement.

**Figure 2.**
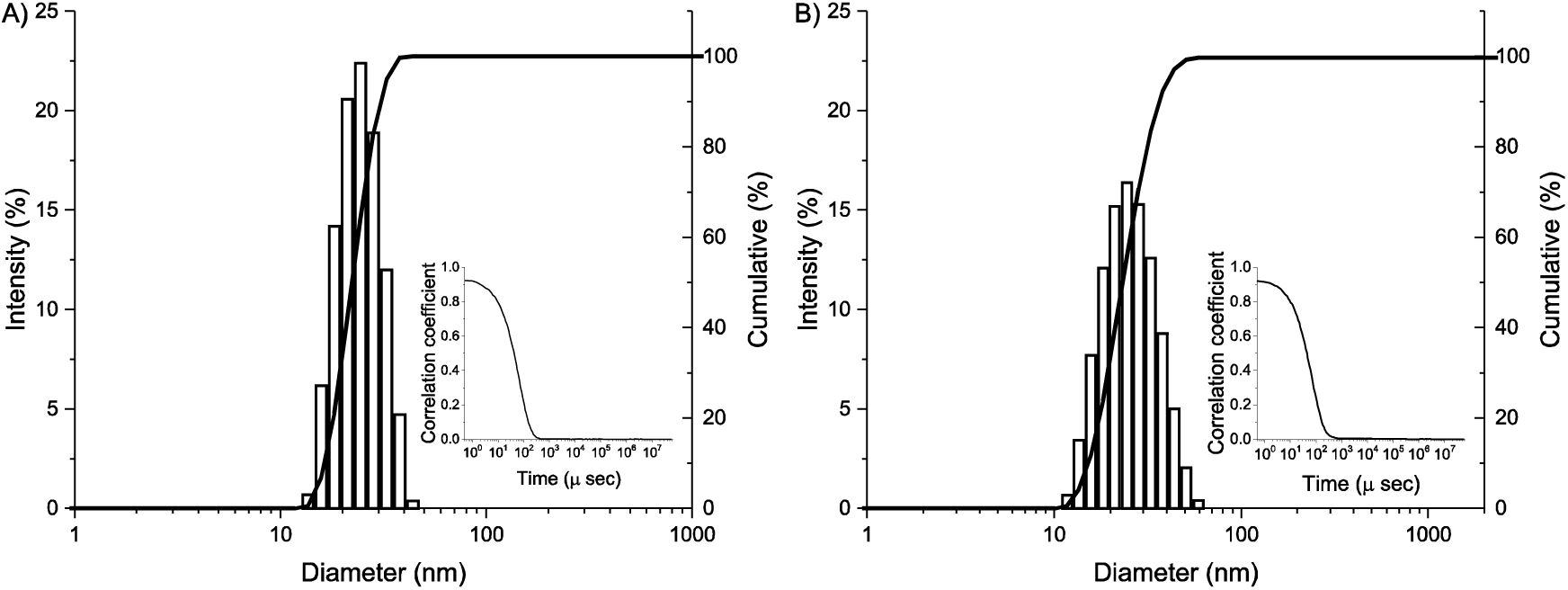
Size distribution of AuNP by PCS. Intensity based distribution and cumulative curves for nanoparticle size distribution for A) citrate-capped AuNP and B) AET-capped AuNP. Inset figures show time decay of correlation.

**Figure 3.**
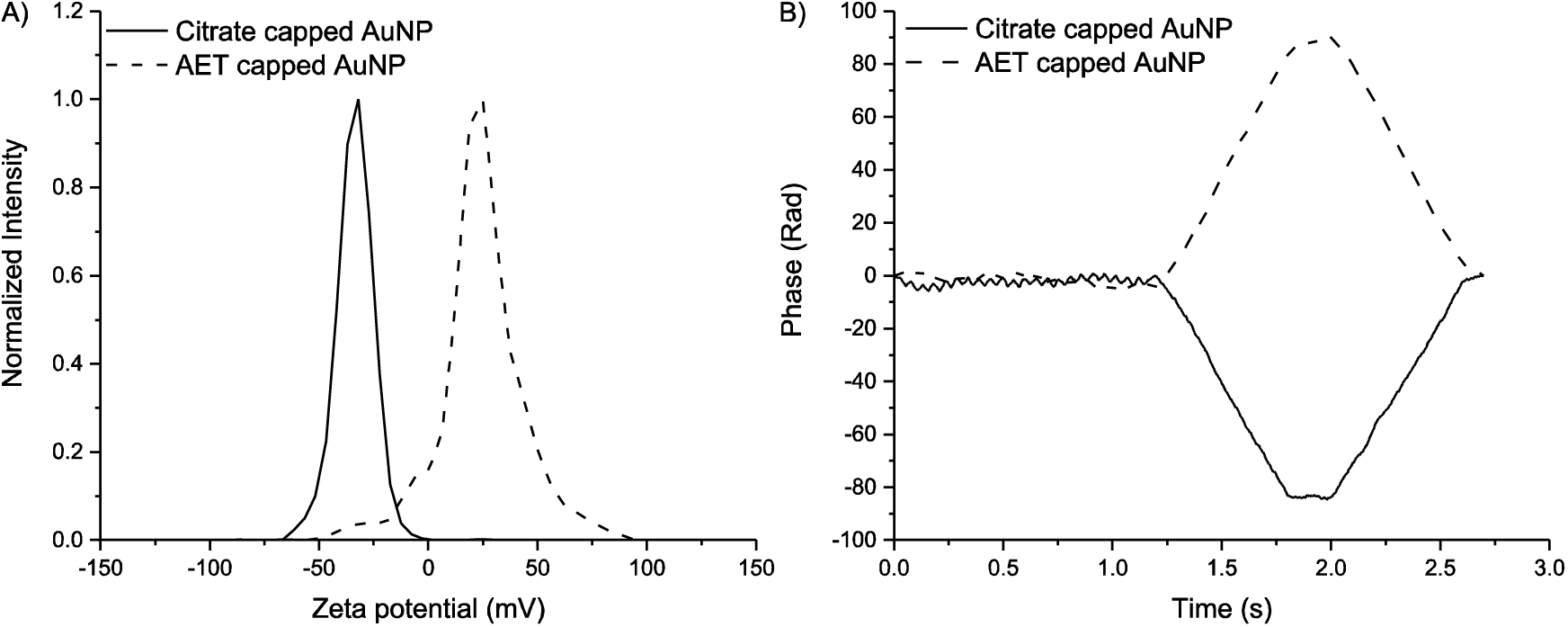
Estimation of zeta potential of AuNP. A) Zeta potential distribution of as prepared citrate-capped and AET-capped AuNP, and B) Phase plots of the same samples.

### Nanoparticle dosing to cells

Before moving ahead with nanoparticle challenge, the conditions of nanoparticle challenge were rationalized. The dosing of nanoparticles is an important but most overlooked aspect of studying the effect of nanoparticle challenge on biological processes *in vitro*. A previously reported theoretical model explaining nanoparticle dosing was based on relationship between concentration of nanoparticles and physicochemical characteristics of nanoparticles such as size, shape, agglomeration state, density etc^25^. Nanoparticles, owing to their small size, undergo Brownian motion when suspended in a solvent and in case of nanobiotechnology, due to high ionic strength, nanoparticles tend to undergo agglomeration. These two phenomena affect the diffusivity and sedimentation rates of nanoparticles in a biological medium thereby influencing the dosing of nanoparticles to the cells *in vitro*. The importance of nanoparticle dosing was studied by Cho E. C. *et al*. wherein they incubated gold nanoparticles with cells either in conventional upright position or in an inverted position^26^. They observed a difference in the uptake of nanoparticles with cells in upright position taking up a significantly higher number of nanoparticles as compared to inverted position.

To rationalize the dose of nanoparticles, it is important to normalize the challenge conditions which can have an influence on dosing to remove the bias in the results. The most important of the conditions are (i) concentration of nanoparticles, (ii) height of media which defines the distance which a nanoparticle needs to travel to interact with the cell, and (iii) agglomeration state of the nanoparticles which defines the size and density of the nanoparticles^25,27^. Moreover, the number of cells can also affect the biological assay. For this work, the number of cells were fixed at 15625 cells/cm^2^. As biological assays are performed in different multiwell plates, the number of cells required to achieve the desired density was calculated and is as shown in **Table 1**. The height of media in a single well is also an important parameter which can influence the results. Therefore, we normalized the volume of media to achieve a height of 0.5 cm in all well plates. For this, different volumes of culture media were suspended in different sized multiwell plates and the height was measured by UV spectrophotometer. **Figure 4** shows the change in height of the media as a function of volume of the media and number of wells in the plate. A linear increase in the height of the media was observed as the volume of media was increased. The linear fit was performed, and all curves had a very high correlation between volume of media and height of media. The volume required for obtaining a height of 0.5 cm was calculated for all well plates and all assays were performed using the same (**Table 2**). For all experiments, the concentration of nanoparticles was estimated by converting molarity of the auric salt used for the synthesis and converting that value into atomic gold. This concentration was estimated to be in µg mL^−1^. All *in vitro* assays were performed using 0.5 cm as height of media, a cellular density of 15625 cells/cm^2^ and a nanoparticle concentration in µg mL^−1^ that lead to a surface area normalized dosing of nanoparticles.

**Table 1.** Number of cells corresponding to a cellular density of 15625 cells/cm^2^.

| Well plate | Growth area (cm<sup>2</sup>) | No. of cells per well |
| --- | --- | --- |
| 96 well | ~ 0.32 | 5,000 |
| 48 well | ~ 0.95 | 15,000 |
| 24 well | ~ 1.90 | 30,000 |
| 12 well | ~ 3.80 | 60,000 |
| 6 well | ~ 9.50 | 150,000 |

**Table 2.** Fit parameters for linear regression performed between height and volume of media in different well plates

| Type of well plate | Slope | Intercept | r<sup>2</sup> | Volume* (μL) |
| --- | --- | --- | --- | --- |
| 96 well | 349.1 | 18 | 0.9995 | 193 |
| 48 well | 1010 | 83.1 | 0.9998 | 588 |
| 24 well | 1914 | 183.8 | 0.9994 | 1141 |
| 12 well | 4016 | 320 | 0.9999 | 2328 |
| 6 well | 9519 | 963.9 | 0.9993 | 5723 |
| *Volume of media required for obtaining a height of 0.5 cm |  |  |  |  |

**Figure 4.**
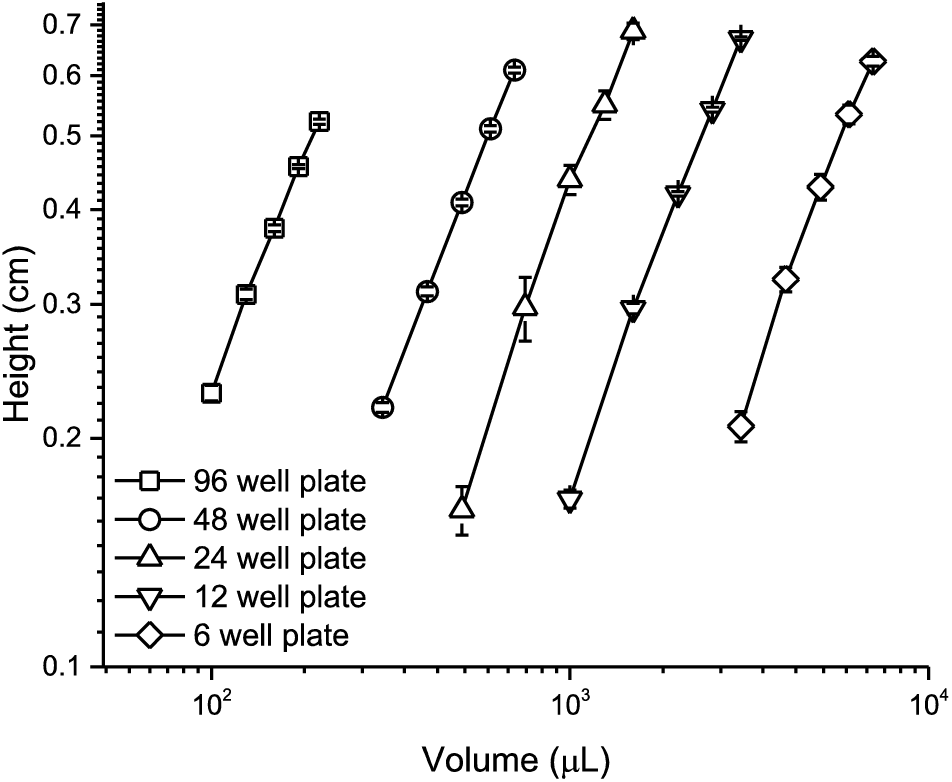
Change in height of media as a function of volume in different well plates used in cell culture (both x and y axes are in logarithmic scale).

### Agglomeration state of AuNP

The agglomeration state of nanoparticles can influence its biological response. For example, agglomeration AuNP has been shown to cause an increase in the cell death^10^. Nanoparticles in their agglomerated state also cause activation of autophagy in cells^11^. Therefore, to make *in vitro* data generated for nanoparticle challenge studies more meaningful, it is important to understand the agglomeration state of nanoparticles in culture media. Nanoparticle concentration, mixing time and time between nanoparticle addition and starting the mixing determine the size of nanoparticle agglomerate formation^28^. Therefore, it is vital to ensure that nanoparticles are properly dispensed in media to minimize agglomeration with time.

To study agglomeration of nanoparticles, complete 2x culture media (DMEM-F12 1:1 mixture supplemented with 10% FBS) was added to equal volume of nanoparticle suspension with immediate mixing of the content using pipette for 60 sec. The nanoparticle suspension in media was then incubated at 37 °C in presence and absence of MDA MB 231 cells. It was observed that adding culture media to nanoparticles causes agglomeration of both anionic and cationic AuNP resulting in an increase of particle size which was indicated by red-shift in the UV-vis absorbance spectra of nanoparticles (**Figure 5**). Further, nanoparticles show agglomeration in presence of cells, however, the red shift observed in case of cationic AuNP was significantly more when compared to anionic AuNP. This can be attributed to the interaction of cationic AuNP with the negative surface charge on MDA MB 231 cells. Similar results have been reported in case of silica nanoparticles, wherein, it was observed that presence of cells changes the dispersion state of nanoparticle suspension^29^. The PCS and LDV analysis to estimate size and zeta potential of these agglomerates could not be performed as the protein content of media was very high and it caused a significant interference (**Figure 6A**). Using centrifugation to separate agglomerates from media was also not successful as it yielded increase in the agglomerate size as indicated by red-shift in UV-vis absorbance spectra of nanoparticles before and after centrifugation (**Figure 6B**). These results indicate that AuNP agglomerate in culture media, however, the physicochemical characteristics of these agglomerates could not be estimated.

**Figure 5.**
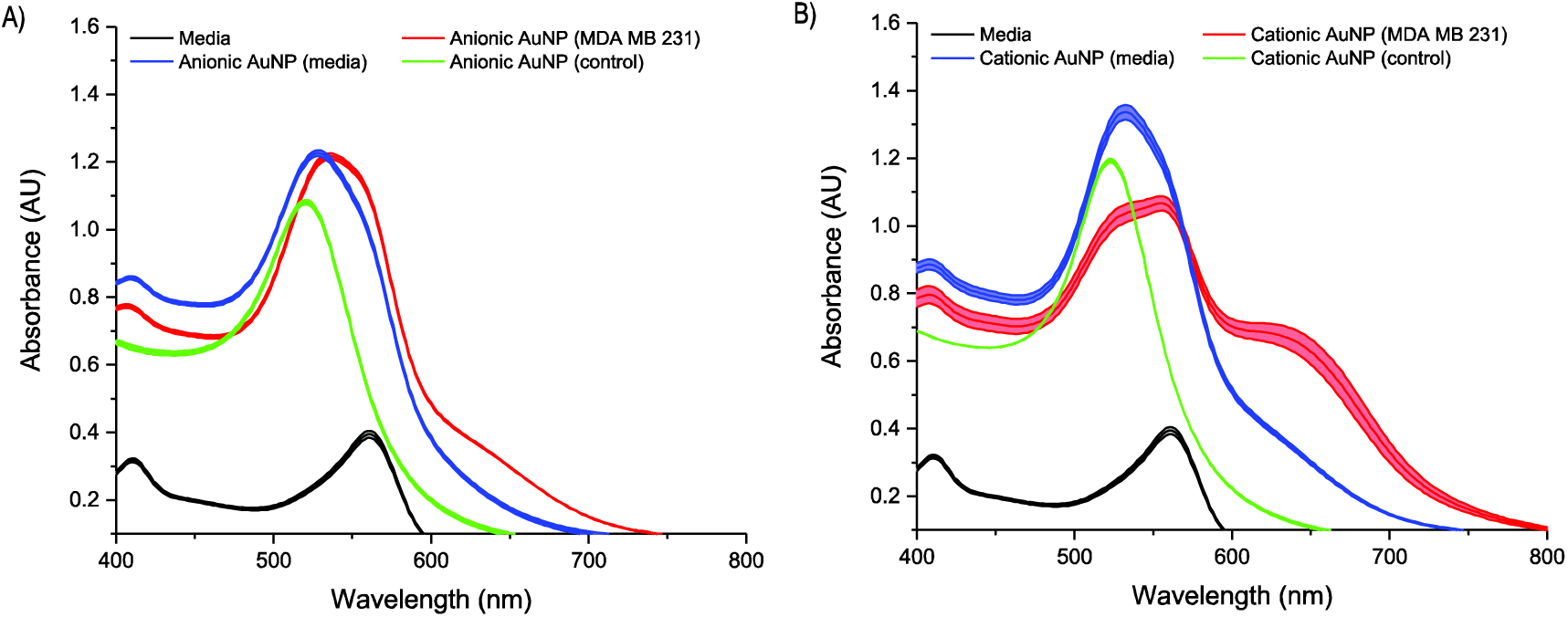
Effect of culture media and cells on nanoparticle agglomeration. UV-vis absorbance spectra of A) anionic AuNP and B) cationic AuNP as a result of exposure to cell culture media and cells. UV-vis absorbance spectrum of media is provided as reference.

**Figure 6.**
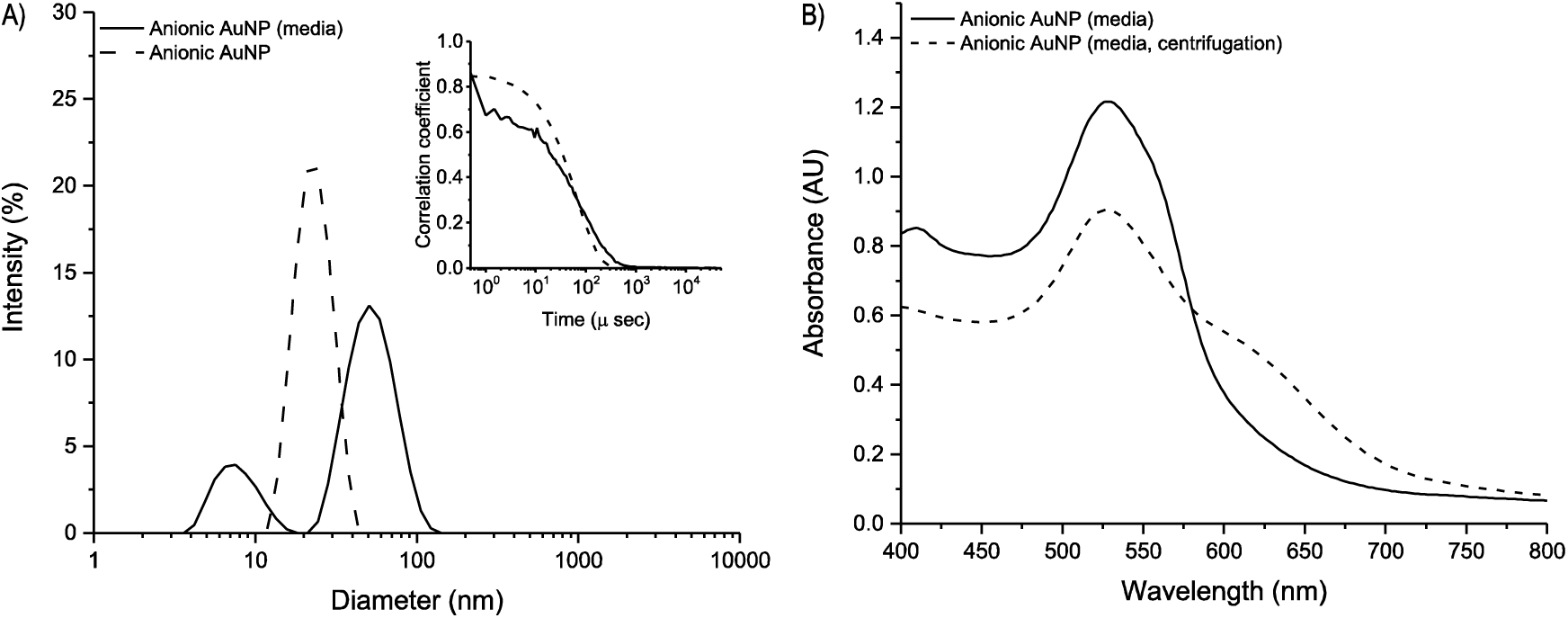
Characterization of nanoparticles after exposure to cell culture media. A) Photon correlation spectroscopy of anionic AuNP before and after incubation with cell culture media. Inset figure shows time decay of correlation, and B) UV-absorption spectra of anionic AuNP after exposure to media and after one cycle of centrifugation to remove cell culture media.

### Effect of AuNP challenge on cell viability

MDA MB 231 and SKBR3 were the two breast cancer cell lines used for the cytotoxicity studies. Cells were seeded at a density of 15,625 cells cm^−2^ in a 96-well plate to study the effect of nanoparticle challenge on cell viability. The nanoparticle suspension prepared in section 0 was used to prepare diluted nanoparticle samples for the study. The resazurin reduction assay was used for quantifying the cell viability after 24 and 48 hours of nanoparticle challenge. Nanoparticle challenge affected the viability of both MDA MB 231 (**Figure 7**) and SKBR 3 (**Figure 8**) in a surface charge and concentration-dependent manner.

**Figure 7.**
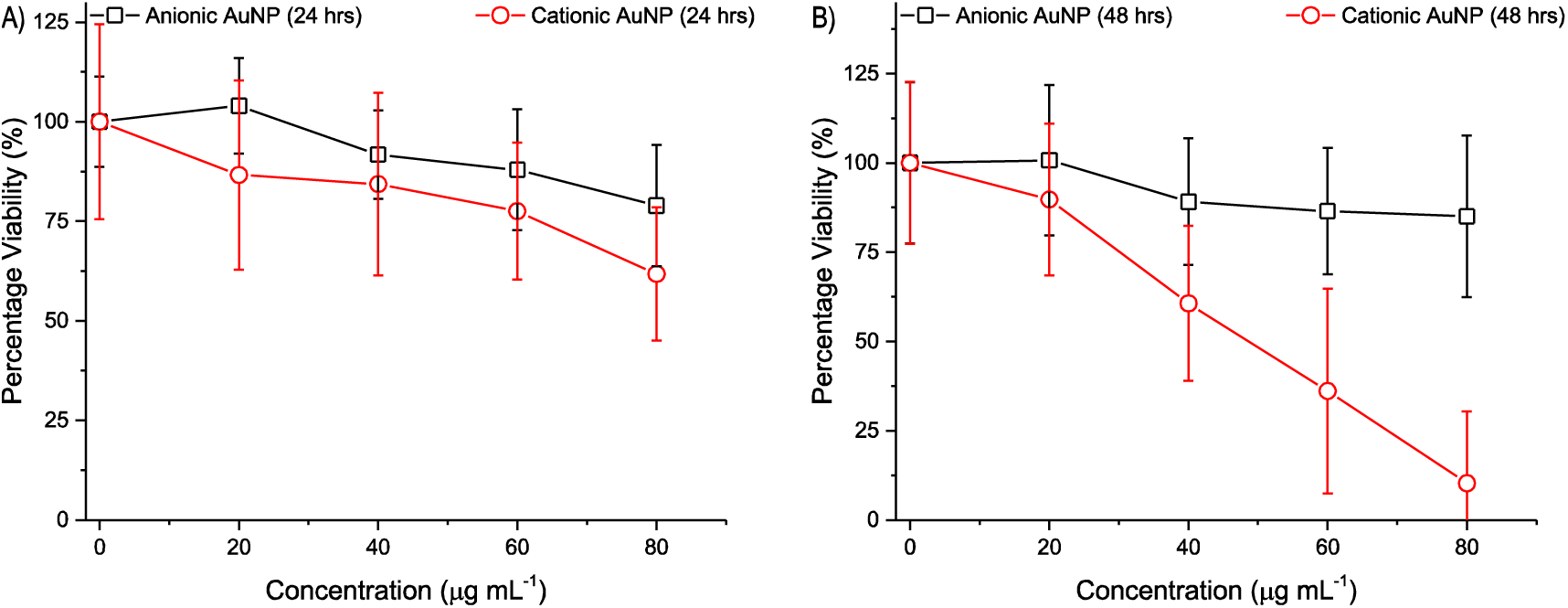
Effect of nanoparticle challenge as a function of concentration on cell viability of MDA MB 231 cell line at A) 24 hours, and B) 48 hours.

**Figure 8.**
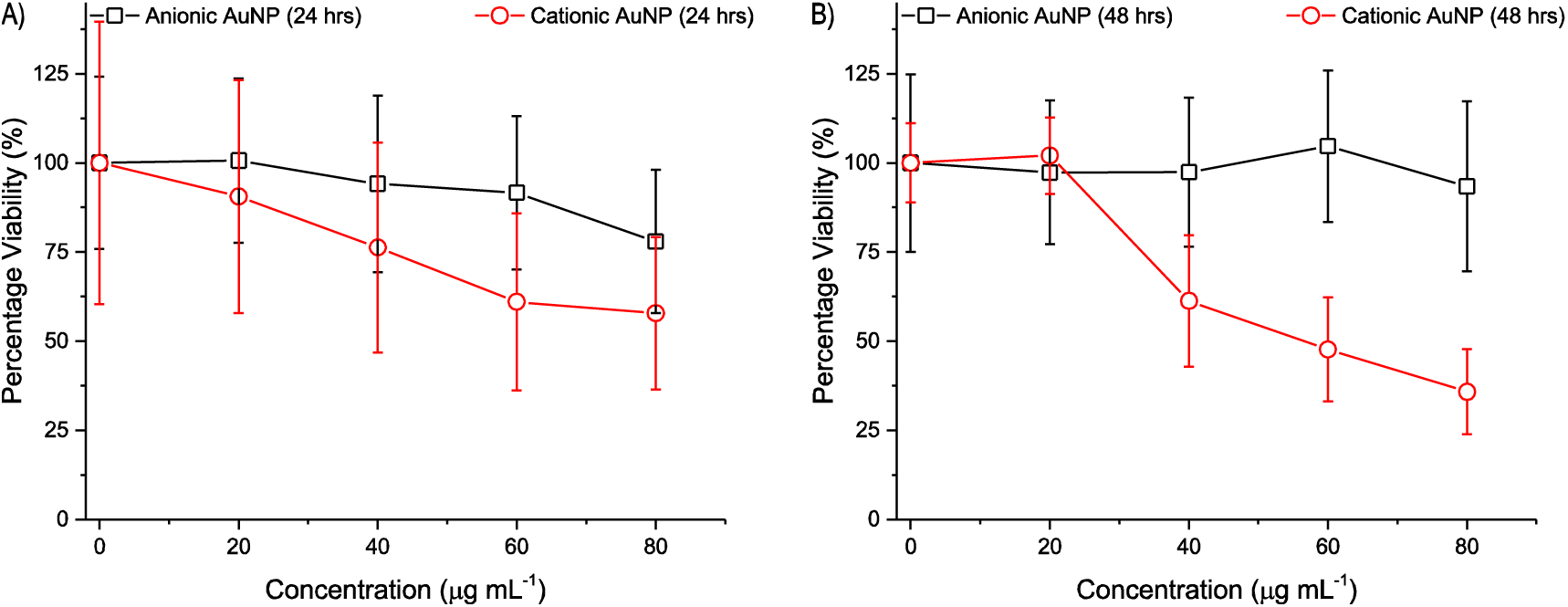
Effect of nanoparticle challenge as a function of concentration on cell viability of SKBR3 cell line at A) 24 hours, and B) 48 hours.

A 3-way Analysis of Variance (ANOVA) analysis was performed to understand the interactions between various variables and their effect on cell viability (**Table 3**). The variability in the MDA MB 231 cytotoxicity profile as a result of nanoparticle challenge (**Figure 7**) was explained majorly by nanoparticle concentration (26.32%) and nanoparticle surface charge density (13.73%). Moreover, there was a significant interaction between nanoparticle concentration and surface charge density (7.034%). Apart from these major contributions, there was a contribution from interactions between nanoparticle concentration, time and surface charge density. On the other hand, in the case of SKBR 3 cell line, the variability of cytotoxicity profile (**Figure 8**) was explained by nanoparticle concentration (19.28%), nanoparticle surface charge density (14.87%), and interaction between the two (9.66%). Further, the contribution from time of exposure and its interaction with nanoparticle concentration, and surface charge density had no significant effect on cell viability.

**Table 3.** Effect of nanoparticle challenge (AuNP concentration, time of exposure, surface charge, and their interactions) on MDA MB 231 and SKBR3 viability (3-way ANOVA, n=6)

| Source of Variation | MDA MB 231 |  | SKBR 3 |  |
| --- | --- | --- | --- | --- |
|  | % of total variation | P value<sup>1</sup> | % of total variation | P value<sup>1</sup> |
| [NP] | 26.32 | **** | 19.28 | **** |
| *t* | 3.837 | ** | 0.03 | ns |
| $\sigma$ | 13.73 | **** | 14.87 | **** |
| [NP] × *t* | 2.847 | ns | 0.33 | ns |
| [NP] × $\sigma$ | 7.034 | ** | 9.662 | ** |
| *t* × $\sigma$ | 3.683 | ** | 1.331 | ns |
| [NP] × *t* × $\sigma$ | 4.197 | * | 2.57 | Ns |
| <sup>1</sup>ns (P > 0.05); * (P ≤ 0.05); ** (P ≤ 0.01); *** (P ≤ 0.001); **** (P ≤ 0.0001)<br>[NP] – concentration of nanoparticle; *t* – time of nanoparticle exposure; $\sigma$ – surface charge of nanoparticles in terms of anionic and cationic | | | | |

The cellular response towards a nanoparticle challenge is pluralistic in nature. Therefore, a 3-way ANOVA analysis was performed to understand the effect of nanoparticle surface charge on cellular viability after nanoparticle challenge. Both MDA MB 231 and SKBR 3 cells did not show a significant difference in their response towards anionic and cationic AuNP (**Table 4**). For anionic AuNP, the concentration of nanoparticle had a significant contribution to the variance of data (9.49%), whereas, all other variables had no significant effect. In the case of cationic AuNP, the nanoparticle concentration (44.68%), time of exposure (5.407%), and their interaction (6.356%) had a significant contribution towards variance in the data. The concentration of nanoparticle was a major determinant of viability in case of cationic AuNP. A 2-way ANOVA analysis of data showed that none of the anionic AuNP concentrations tested had any significant effect on cell viability of either cell types (**Table 5**). Whereas, cationic AuNP caused cytotoxicity in both concentration and time-dependent manner for both cell types. Therefore, AuNP caused cytotoxicity in a time and concentration-dependent manner with nanoparticle surface charge playing a significant role. Moreover, there was no significant difference in the response of MDA MB 231 and SKBR 3 cell lines towards nanoparticle challenge. All future studies were thus limited to MDA MB 231 cell line.

**Table 4.** Effect of nanoparticle surface charge on the viability of MDA MB 231 and SKBR3 cells (3-way ANOVA, n=6)

| Source of Variation | Anionic AuNP |  | Cationic AuNP |  |
| --- | --- | --- | --- | --- |
|  | % of total variation | P value<sup>1</sup> | % of total variation | P value<sup>1</sup> |
| [NP] | 9.49 | * | 44.68 | **** |
| Cell line | 0.7037 | ns | 0.1502 | ns |
| t | 0.4699 | ns | 5.407 | *** |
| [NP] × Cell line | 1.508 | ns | 0.7922 | ns |
| [NP] × t | 1.58 | ns | 6.356 | ** |
| Cell line × t | 0.5539 | ns | 1.295 | ns |
| [NP] × Cell line × t | 0.493 | ns | 0.7991 | Ns |
| <sup>1</sup>ns (P > 0.05); * (P ≤ 0.05); ** (P ≤ 0.01); *** (P ≤ 0.001); **** (P ≤ 0.0001)<br>[NP] – concentration of nanoparticle; Cell line – cell type (either MDA MB 231 or SKNR3);<br>t – time of nanoparticle exposure |  |  |  |  |

**Table 5.**
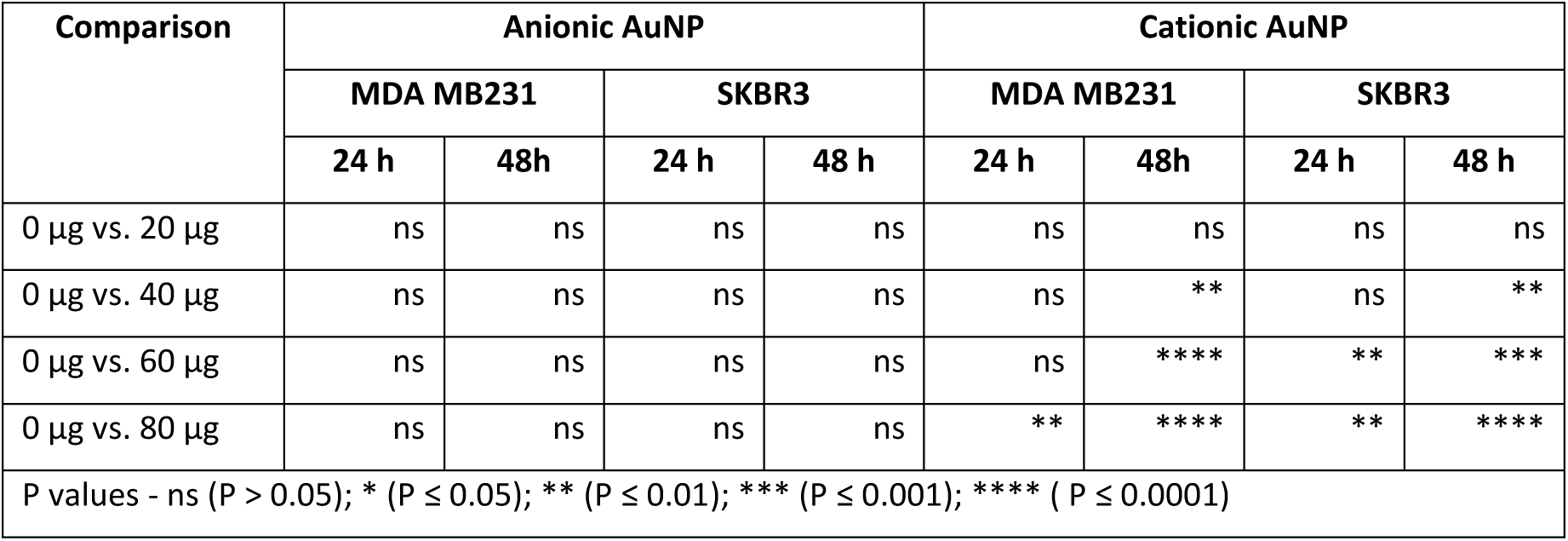
Effect of concentration of nanoparticles on cell viability of MDA MB 231 and SKBR3 cells (2-way ANOVA)

### Effect of AuNP challenge on plasma membrane integrity

The plasma membrane is a double-layered lipid membrane which encapsulates the cytoplasm of eukaryotic cells. Nanoparticles covered with a biomolecular corona interact with the plasma membrane of cells and are actively internalized by cells. Nanoparticles are internalized at the endocytic hot spots present at the apical surface of the cells. Endocytic hot spots are generally rich in cholesterol and form raft like structures in the plasma membrane. It has been reported that AuNP are actively taken up by the cells at their apical surface only^14^. Nanoparticle internalization is associated with the uptake of a large amount of plasma membrane. The resulting plasma membrane damage might initiate an instantaneous repair mechanism mediated by intracellular calcium concentration^30^ leading to the plasma membrane replenishment by exocytosis of lysosomes^31^. Therefore, nanoparticle uptake should be associated with exocytosis if cells are internalizing excess of plasma membrane. It has been observed that cells do show exocytosis of nanoparticles^32^ indicating that nanoparticle uptake results in enhanced plasma membrane internalization. A mismatch in the turnover of plasma membrane can affect the plasma membrane structure and consequently its physicochemical characteristics. Therefore, we hypothesized that, ***nanoparticle interaction with the plasma membrane and their subsequent internalization by the cells could cause changes in the physicochemical characteristics of plasma membrane***. To test this hypothesis, the nanoparticle-mediated changes in the physicochemical characteristics of plasma membrane including integrity, fluidity, and surface charge of the plasma membrane were studied.

Physicochemical characteristics of plasma membrane play an important role in sensing of environmental stress and signal transduction. Therefore, the interaction between biomolecular corona-coated nanoparticles and plasma membrane could be an important determinant for nanoparticle-cell interaction and ensuing cellular response. FDA/PI staining was used to study the effect of nanoparticle challenge on plasma membrane integrity as a function of nanoparticle concentration and time of exposure. FDA is a vital dye which stains live cells green, whereas, PI is a plasma membrane impermeable nuclear stain which stains nucleus red as a result of the loss of integrity of the plasma membrane. The MDA MB 231 cells were treated with anionic and cationic AuNP and plasma membrane integrity was assessed by FDA/PI staining after 3 and 12 hours. There was no effect observed for anionic AuNP challenge at both low and high doses. However, cationic AuNP had a concentration and time-dependent effect on plasma membrane integrity at 12 hours. Cationic AuNP at a concentration of 80 µg mL^−1^ caused a significant plasma membrane damage at 12 hours which lead to a reduction in the number of cells per field (**Figure 9**). these results are in line with previous reports, wherein, cationic polystyrene nanoparticles have been shown to elicit a cytotoxic response by compromising the integrity of plasma membrane^33^.

**Figure 9.**
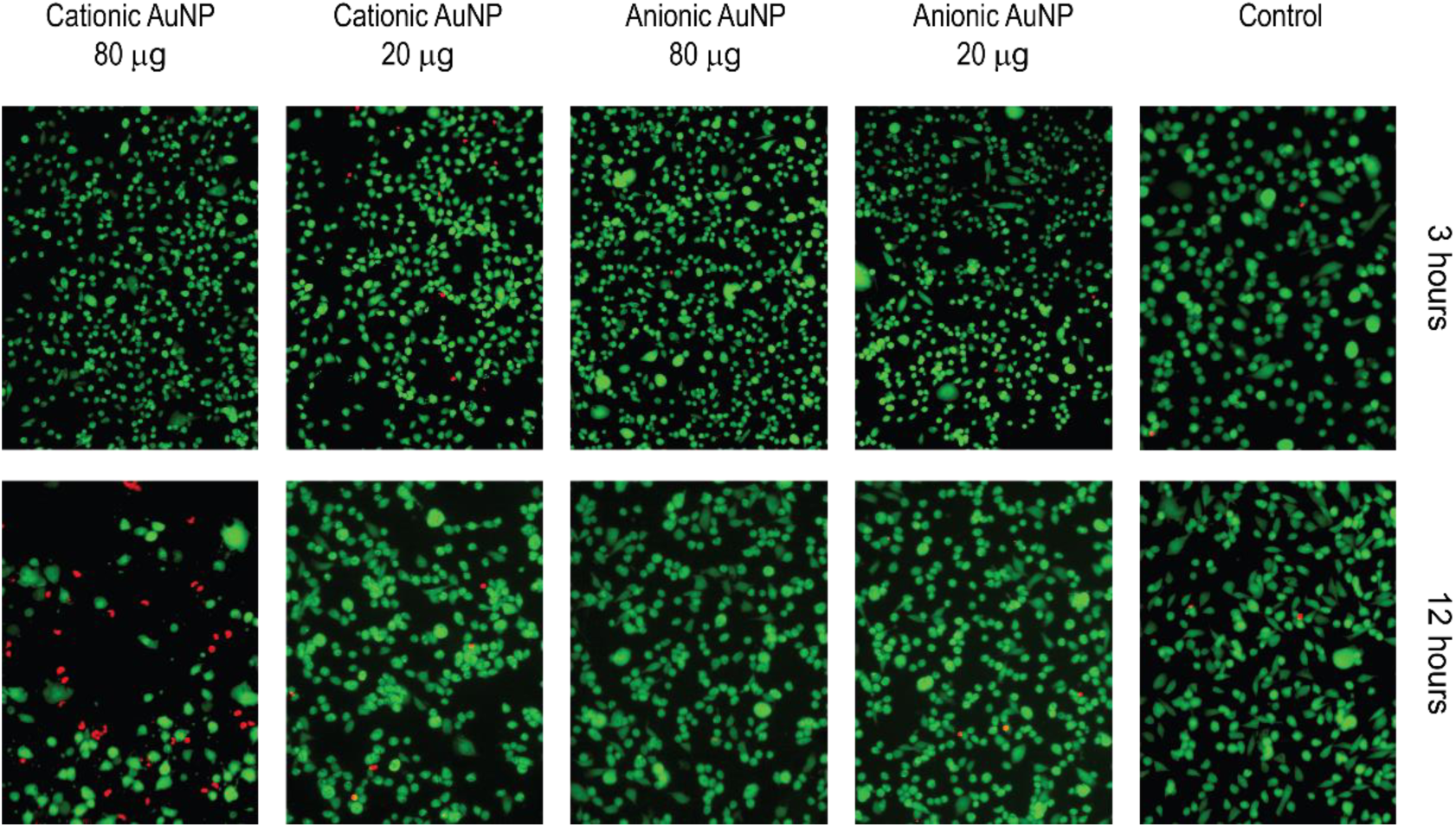
Plasma membrane integrity of MDA MB 231 cells after 3 hours and 12 hours of exposure to anionic and cationic AuNP. Green staining is due to live cells and red staining is due to nuclear staining of cells with compromised cell membrane.

### Effect of AuNP challenge on cell surface charge

The plasma membrane of mammalian cells is a bilayer structure formed by phospholipids with an overall negative surface charge. The phospholipids are distributed asymmetrically in the phospholipid bilayer with the outer leaflet comprising majorly of choline-containing phospholipids while the cytosolic leaflet comprises majorly of aminophospholipids. The asymmetry of charged phospholipids causes counterion accumulation around the plasma membrane giving rise to a transmembrane potential. The transmembrane potential is instrumental in ion transport, especially Ca^2+^ transport across the plasma membrane and is known to control many cellular processes^34^.

Charge based electrostatic interactions also play a significant role in defining the nanoparticle-bio interface. Nanoparticles interact with the plasma membrane of cells in a charge dependent manner. Positively charged nanoparticles tend to interact more with cells as compared to negatively charged nanoparticles^35^. It has been previously reported that nanoparticle interaction with cells causes a change in surface charge density of cells. For example, anionic nanoparticle uptake by MCF 7 (breast cancer cells) and MCF 10A (normal breast cells) caused a change in the surface charge of these cells^15^. On incubation with anionic iron oxide nanoparticles, there was a decrease in the zeta potential of MCF 10A cells, whereas, the zeta potential of MCF 7 cells increased. Any change in the surface charge density can potentially alter the transmembrane potential of the plasma membrane as it alters the accumulation of counterions on the outer leaflet of the plasma membrane. It has also been shown that nanoparticle internalization by cells can cause a change in the transmembrane potential as a function of nanoparticle surface charge. Arvizo *et al.* Cationic gold nanoparticles cause plasma membrane depolarization in mammalian cells whereas, anionic nanoparticles do not have any effect on transmembrane potential^7^. Therefore, ***it is imperative to understand the effect of nanoparticle challenge on cell surface charge.***

MDA MB 231 cells were incubated with low (20 µg mL-1) and high (80 µg mL-1) concentration of anionic and cationic AuNP for 3 and 6 hours to study the effect of AuNP challenge on cell surface charge. A 6 hours’ incubation time was selected as 12 hours of incubation with cationic AuNP resulted in cell death (section 0). The culture media was removed after incubation, and cells were washed with PBS and were detached using a non-enzymatic cell dissociation reagent to avoid lysis of cell surface proteins by enzymes. Some previous studies have used harsh enzymatic treatments, such as trypsin-EDTA, to detach cells before measuring surface charge. Also, the time required for the detachment of cells from the substrate is variable, and it ranges from 3-5 minutes for MCF 7 cells and 15 minutes for MCF 10A and therefore introduces a bias in the results^15^. The detached cells were then centrifuged and resuspended in a solution of known ionic strength to measure the zeta potential.

The zeta potential is a function of the ionic strength of the media as ions present in the medium affect the electrophoretic mobility of cells. The extracellular medium has an osmolality of 380 mOs. Zeta potential of MDA MB 231 cells was measured as a function of extracellular ionic strength while keeping the osmolality of the media constant by adding glucose to the medium. Zeta potential of cells changed with the increase in conductivity and the value of zeta potential stabilized above 60 mM NaCl concentration (**Figure 10A**). However, the phase plots of Laser Doppler Velocimetry (**Figure 10B**) started showing charging of solution beyond 38.8 mM NaCl. Beyond 53.2 mM NaCl, the curve was unstable in the fast-field reversal region (below 1.0 second) while there was no measurement recorded in the slow-field reversal region (1.0-2.5 seconds). Therefore, the equipment failed to estimate the zeta potential at a higher ionic strength of extracellular medium (**Figure 10C**). All the zeta potential values were, therefore, measured using 10 mM of NaCl while maintaining the overall osmolality of the extracellular media at 380 mOs. MDA MB 231 cells showed a zeta potential of around -20 mV at 3 and 6 hours time points (**Figure 10D**). Incubation of cells with low concentration (20 µg mL^−1^) of anionic AuNP did not cause any significant change in zeta potential values at both time points. However, high concentration (80 µg mL^−1^) of anionic AuNP caused a significant depolarization at 3 hours, but the cells retained their basal surface charge after 6 hours of incubation. An effect similar to high anionic AuNP concentration was observed for low concentration (20 µg mL^−1^) of cationic AuNP. In the case of higher concentration of cationic AuNP, the cells showed a significant depolarization which was persistent up to 6 hours. Therefore, nanoparticles showed a surface charge dependent effect on the cell surface charge density of MDA MB 231 cells.

**Figure 10.**
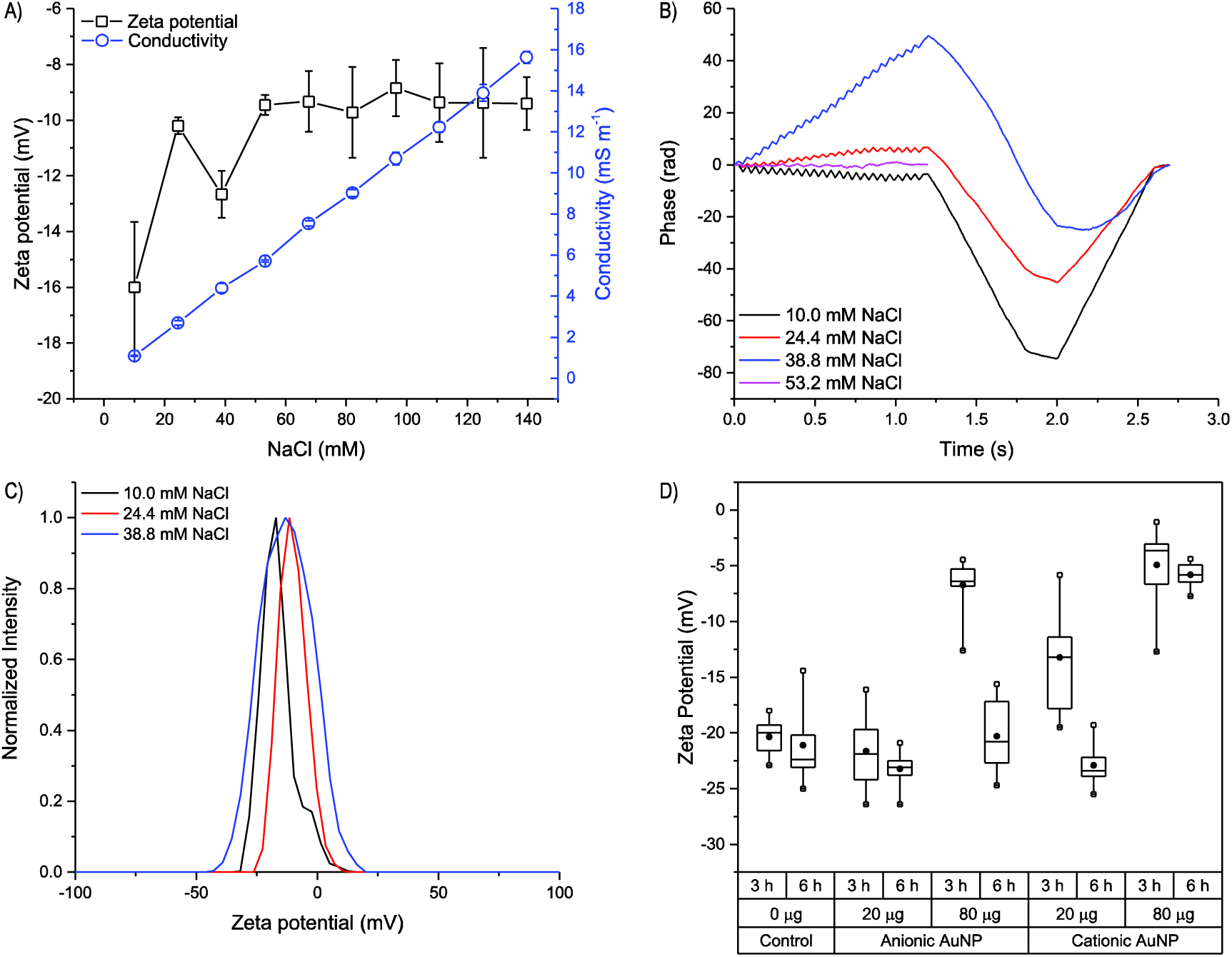
Effect of nanoparticle challenge on surface charge of MDA MB 231 cells. A) Change in zeta potential of MDA MB 231 cells as a result of change in ionic strength of the media. Changes in the B) Phase plot and C) Zeta potential distribution of MDA MB 231.cells D) Changes in the zeta potential of MDA MB 231 cells as a result of challenge by anionic and cationic AuNP after 3 and 6 hours.

### Effect of AuNP challenge on plasma membrane fluidity

Plasma membrane plays an important role as an environmental stress sensor. Cold temperatures cause membrane rigidity, and higher temperatures cause membrane fluidity^36^. Plasma membrane hyperfluidization during heat shock causes activation of heat stress response, and therefore, plasma membrane fluidity acts as a heat sensor^16,36,37^. Therefore, any change in plasma membrane fluidity as a result of nanoparticle challenge can potentially have the same effect on cells as caused by environmental stress.

Pyrene excimerization was studied to elucidate the effect of nanoparticle challenge on plasma membrane microviscocity of MDA MB 231 cells. Pyrene, a polycyclic organic compound which shows excimerization, was used for measuring membrane microviscosity. Excimerization is a phenomenon in which a molecule in excited state can interact with a ground state molecule to form an excited dimer which is known as excimer. The rate of excimer formation is diffusion dependent and thus is related to the local viscosity of the surrounding medium^22^. Further, to test the feasibility of this technique, the plasma membrane fluidity of MDA MB 231 cells was altered using benzyl alcohol treatment. Benzyl alcohol causes plasma membrane fluidization^38^. A decrease in the amount of free pyrene in the surrounding medium with an increase in benzyl alcohol concentration was indicative of an increase in the pyrene excimerization and, therefore, fluidity of the plasma membrane (**Figure 11A**). Further, to study the effect of nanoparticle treatment on plasma membrane fluidity of MDA MB 231 cells, these cells were treated with different concentrations of anionic and cationic AuNP for 3 hours. Low levels of both anionic and cationic AuNP did not change plasma membrane fluidity (**Figure 11B**). However, higher concentrations of cationic AuNP increased the fluidity of the plasma membrane, whereas, anionic AuNP caused a slight increase in the rigidity of plasma membrane at higher concentrations.

**Figure 11.**
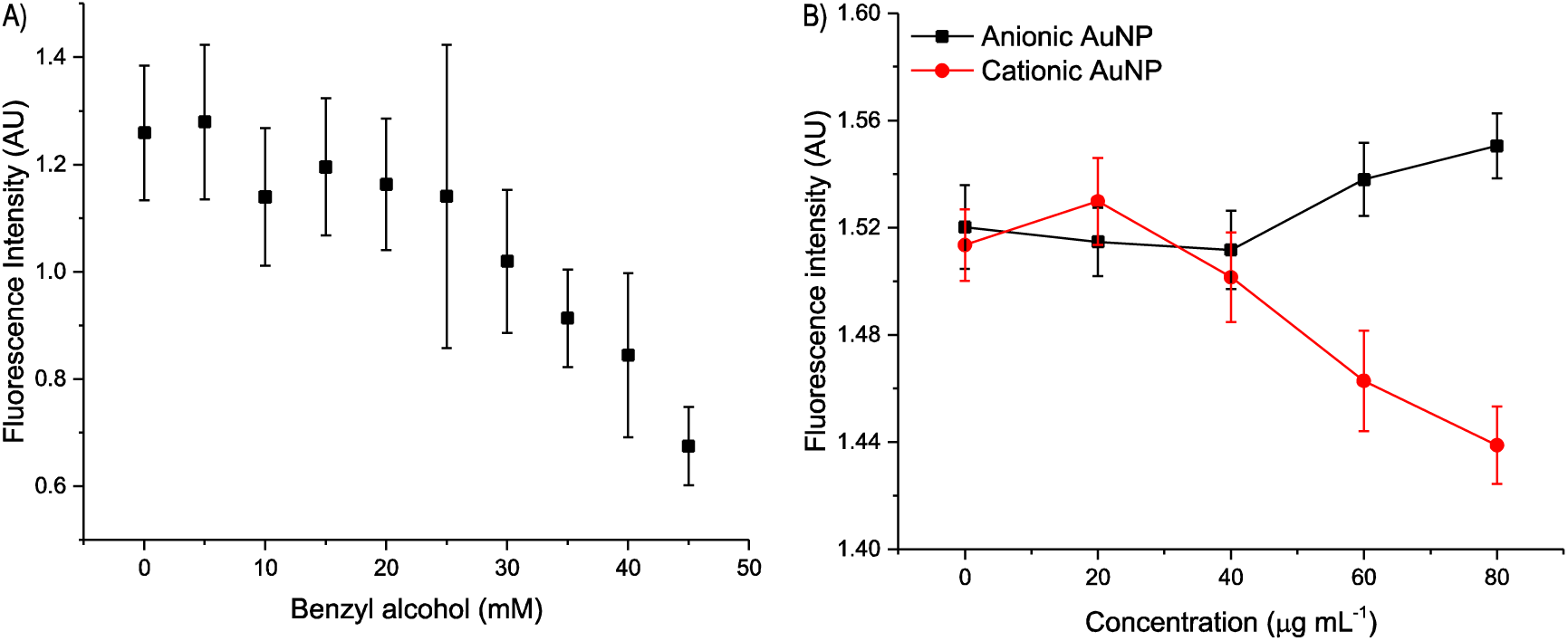
Changes in the plasma membrane fluidity of MDA MB 231 cells as a result of A) Benzyl alcohol treatment, and B) Three hours of challenge with anionic and cationic AuNP.

These results are consistent with the results obtained for a single phospholipid bilayer incubated with the nanoparticles, wherein, anionic nanoparticles caused local gelation, and cationic nanoparticles caused local fluidization^39^. Moreover, the changes in plasma membrane fluidity can also explain the heat shock mediated toxic response of AuNP as observed in *Drosophila*^40^. Plasma membrane fluidization is known to activate phospholipase C which in turn increases the concentration of inositol-1,4,5-triphosphate (IP3) and diacylglycerol inside the cells^41^. IP3 increases the intracellular Ca^2+^ concentration by activating IP3 receptors present on the endoplasmic reticulum^16^. As stated earlier, cationic AuNP challenge also results in similar intracellular Ca^2+^ spikes^7^. Therefore, we hypothesized that ***nanoparticle-mediated changes in the fluidity of the plasma membrane could potentially influence the ensuing cellular responses***. The heat shock is also associated with an increase in the cytosolic concentration of reactive oxygen species and is also known to cause apoptosis by activation of mitochondrial pathway mediated by caspase 9^42^. ***Hence, nanoparticle-mediated changes in the physicochemical characteristics of the plasma membrane, especially plasma membrane fluidity, could influence the mitochondrial bioactivity and cytosolic ROS levels.***

### Effect of AuNP challenge on mitochondrial bioactivity and cytosolic reactive oxygen species (ROS) concentration

Nanoparticle-mediated cytotoxicity is known to be a consequence of intracellular oxidative stress generated by nanoparticle internalization^43^. Redox homeostasis controls cellular processes at the level of transcription (NF-*κ*B, AP-1, Nrf2, HIF), signal transduction (MAPK, PI3k/Akt) and cell viability (caspase, Bcl-2, cytochrome c, p53). Cells respond to different levels of oxidative stress in a hierarchical manner^44^. The first level of response involves activation of antioxidant response element and a cytoprotective response. At intermediate levels of ROS, MAPK pathway and NF-κB pathway get activated which cause cellular proliferation and cytokine release respectively. Higher levels of ROS are detrimental to cell and it causes cell death by apoptosis or necrosis. Intestinal cells have been shown to have a concentration-dependent response to hydrogen peroxide challenge (oxidative stress) *in vitro*. These cells showed proliferation at a mild stress (<10 µM L^−1^), apoptosis at an intermediate stress (10-50 µM L^−1^) and necrosis at high stress (>100 µM L^−1^)^45^. It is known that mitochondria play a significant role in the oxidative state of the cell as it is the primary source of intracellular ROS in eukaryotic cells^46^. Hence, mitochondrial ROS can contribute to the overall increase in cytosolic ROS concentration after the nanoparticle challenge.

Apart from mitochondria, NADPH oxidase (NOX) also plays a role in the generation of ROS. NOX actively forms superoxide ions which get converted into hydrogen peroxide or peroxynitrite by cytosolic enzymes. Mild hyperthermia causes NOX-mediated activation of hypoxia-inducible factor 1 (HIF 1) in MDA MB 231 cells followed by activation of extracellular signal-regulated kinases (ERK) pathway^47^. As plasma membrane hyperfluidization acts as a sensor for hyperthermia, therefore we hypothesized that ***any process leading to membrane fluidization can potentially cause activation of NOX and a significant increase in the cytosolic*** ROS levels. Further, NOX-mediated ROS induces the activation of ATP-dependent potassium channels present in mitochondrial membrane^26,48^<u>_ENREF_118</u>. The activation of potassium channels results in loss of mitochondrial membrane potential and release of mitochondrial ROS, eventually causing an elevated level of oxidative stress. This interplay between NOX-derived and mitochondrial ROS could have an implication in oxidative stress-mediated cellular responses caused by nanoparticle challenge. Previous reports have demonstrated both intermediate and high-level ROS response by different cells as a result of AuNP challenge. For example, oxidative stress is the primary mechanism behind AuNP cytotoxicity in HL60 and HepG2 cell lines^49^. Treatment with an antioxidant (N-acetyl cysteine) showed a cytoprotective effect, whereas, depletion of glutathione (GSH) and superoxide dismutase (SOD) caused an elevation in the ROS levels which together confirmed the role of oxidative stress in cell death. Moreover, AuNP interaction with cancer cells also results in abrogation of MAPK pathway^50^, whereas, their interaction with primary cells causes activation of MAPK pathway^51^. As noted earlier, intermediate level of oxidative stress activates MAPK. Collectively, these results indicate that oxidative stress has a significant role in determining the cellular response after nanoparticle challenge.

The mitochondrial membrane potential (MMP) and cytosolic ROS levels were measured to study the effect exposure (3 hours incubation) of anionic and cationic AuNP on MDA MB 231 cells MMP was estimated by mitochondrial accumulation of rhodamine 123 (Rh 123). Rh123 is a lipophilic, cell-permeable cationic dye which accumulates inside the mitochondria as a function of MMP, and hence, mitochondrial depolarization results in the loss of Rh123 fluorescence. A significant change in the MMP of MDA MB 231 cells was observed for both anionic and cationic AuNP (**Figure 12A**). The concentration of the nanoparticles was the primary source of variation observed in the data (**Table 6**). Anionic and cationic AuNP caused a significant change in MMP above 20 µg mL^−1^ (**Table 7**). Moreover, the surface charge of nanoparticles had no significant effect on MMP at all concentrations of nanoparticles tested (**Table 8**). These results indicated that nanoparticle interaction with the cells caused a change in the MMP irrespective of the surface charge of nanoparticles. Citrate-capped AuNP of 13 nm have also been shown to cause a similar effect on the MMP of rabbit articular chondrocytes^52^.

**Table 6.** Effect of nanoparticle challenge on mitochondrial membrane potential (MMP) and reactive oxygen species (ROS) levels of MDA MB 231 cells (2-way ANOVA, n=6)

| Source of variation | MMP |  | ROS |  |
| --- | --- | --- | --- | --- |
|  | % of variation | P value<sup>1</sup> | % of variation | P value<sup>1</sup> |
| [NP] | 54.31 | **** | 12.38 | ** |
| $\sigma$ | 2.784 | ns | 43.77 | **** |
| [NP] × $\sigma$ | 3.851 | ns | 12.67 | ** |

<sup>1</sup>ns (P > 0.05); \* (P ≤ 0.05); \*\* (P ≤ 0.01); \*\*\* (P ≤ 0.001); \*\*\*\* (P ≤ 0.0001)
[NP] – concentration of nanoparticle; $\sigma$ – surface charge of nanoparticles in terms of anionic and cationic

**Table 7.** Effect of concentration of nanoparticle on mitochondrial membrane potential (MMP) and reactive oxygen species (ROS) levels of MDA MB 231 cells (2-way ANOVA, n=6)

| Comparison | Anionic AuNP |  | Cationic AuNP |  |
| --- | --- | --- | --- | --- |
|  | MMP | ROS | MMP | ROS |
| 0 µg vs. 20 µg | ns | ns | ns | * |
| 0 µg vs. 40 µg | ** | ns | ** | **** |
| 0 µg vs. 60 µg | *** | ns | **** | **** |
| 0 µg vs. 80 µg | * | ns | **** | ** |

P value - ns (P > 0.05); \* (P ≤ 0.05); \*\* (P ≤ 0.01); \*\*\* (P ≤ 0.001); \*\*\*\* (P ≤ 0.0001)

**Table 8.** Effect of nanoparticle surface charge on mitochondrial membrane potential (MMP) and reactive oxygen species (ROS) levels of MDA MB 231 cells (2-way ANOVA, n=6)

| Anionic AuNP vs. Cationic AuNP | MMP | ROS |
| --- | --- | --- |
| 0 µg | ns | ns |
| 20 µg | ns | *** |
| 40 µg | ns | **** |
| 60 µg | ns | **** |
| 80 µg | ns | ** |
| P value - ns (P > 0.05); * (P ≤ 0.05); ** (P ≤ 0.01); *** (P ≤ 0.001); **** (P ≤ 0.0001) |  |  |

**Figure 12.**
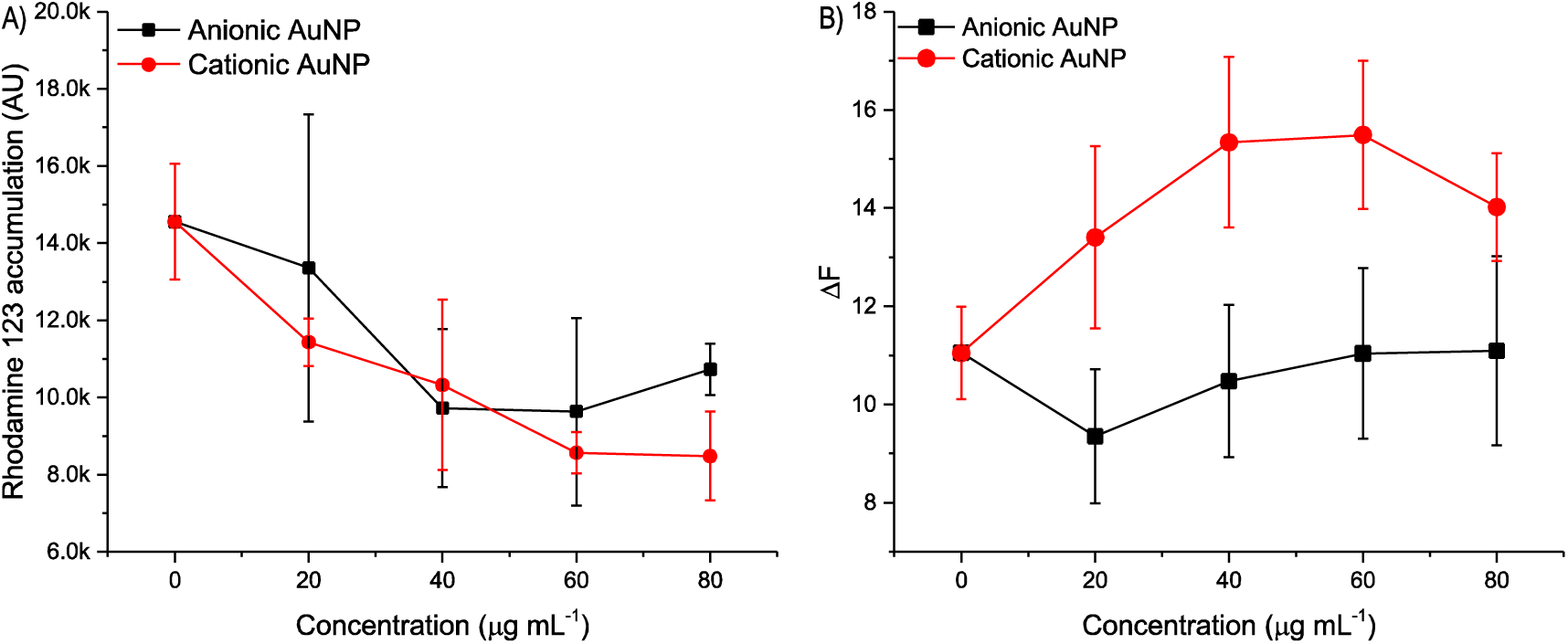
Effect of nanoparticle challenge on A) Rhodamine 123 accumulation in mitochondria, and B) Reactive oxygen species level in MDA MB 231 cells

Oxidative stress generated by nanoparticle challenge in MDA MB 231 cells was estimated by DCF-DA assay, wherein, non-fluorescent and cell membrane permeable DCF-DA is reduced to a fluorescent and cell membrane impermeable product by the action of intracellular hydrogen peroxide. The ratio of the change in fluorescence (at 30 and 120 minutes) after nanoparticle challenge is an indicator of the levels of cytosolic ROS. It was observed that cationic AuNP caused an increase in overall oxidative stress, whereas, anionic AuNP showed no such effect (**Figure 12B**). The oxidative stress generated by AuNP challenge was significantly affected by the surface charge of nanoparticles, the concentration of nanoparticles and due to the interaction between these two parameters (**Table 6**). Cationic AuNP challenge caused a significant increase in the oxidative stress at all concentrations tested, whereas, for anionic AuNP, there was no significant change in the levels of ROS (**Table 7**). Moreover, at all concentrations tested, the oxidative stress generated by cationic AuNP was significantly higher when compared with anionic AuNP (**Table 8**). These results highlight the role of oxidative stress in governing the cytotoxic response of AuNP challenge; however, the role of mitochondrial ROS in determining the cytotoxic response is questionable.

### Principal Component Analysis of AuNP mediated cytotoxicity

PCA is used to reduce the number of factors explaining the variation in the data. Principal factors identified are linear combinations of factors used for the analysis. PCA can, therefore, provide the information about the relative importance of different factors in obtaining a particular outcome. Therefore, a PCA was performed to estimate the contribution of various factors (which are, cell viability, nanoparticle concentration, MMP, plasma membrane rigidity, and cytosolic ROS concentration) towards cytotoxic response of nanoparticles. For PCA analysis, all values were transformed to the percentage points with respect to the untreated control group. Since AuNP surface charge is not a continuous variable, therefore, it was used as a supplementary variable in the PCA analysis. Correlation matrix of the data based on Pearson’s correlation coefficient is provided in **Table 9**. The correlations which were significantly different than 0 at a significance level of α=0.05 are in boldface.

**Table 9.**
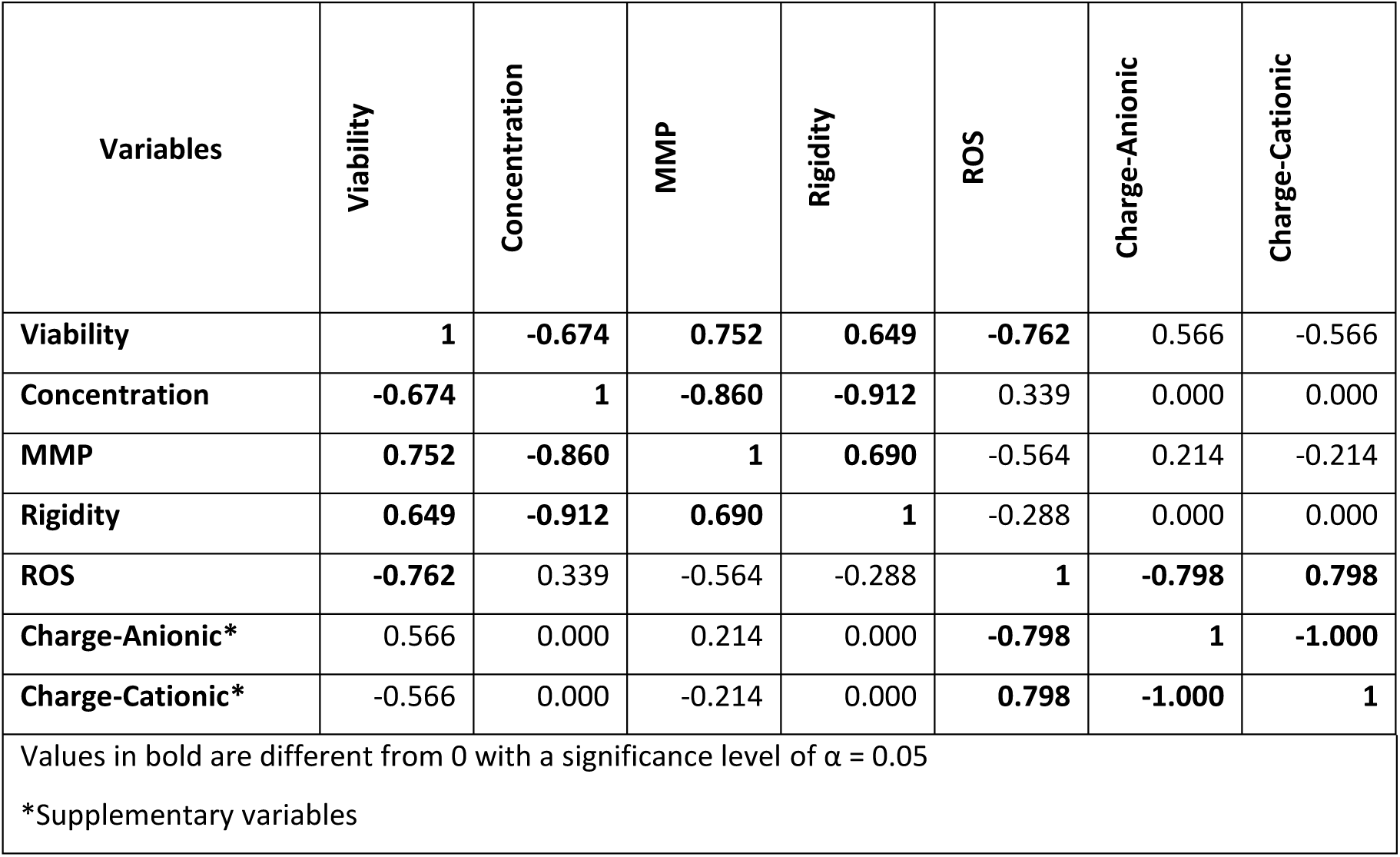
Correlation matrix (Pearson’s correlation coefficient) for the PCA analysis.

| Variables | Viability | Concentration | MMP | Rigidity | ROS | Charge-Anionic | Charge-Cationic |
| --- | --- | --- | --- | --- | --- | --- | --- |
| Viability | 1 | -0.674 | 0.752 | 0.649 | -0.762 | 0.566 | -0.566 |
| Concentration | -0.674 | 1 | -0.860 | -0.912 | 0.339 | 0.000 | 0.000 |
| MMP | 0.752 | -0.860 | 1 | 0.690 | -0.564 | 0.214 | -0.214 |
| Rigidity | 0.649 | -0.912 | 0.690 | 1 | -0.288 | 0.000 | 0.000 |
| ROS | -0.762 | 0.339 | -0.564 | -0.288 | 1 | -0.798 | 0.798 |
| Charge-Anionic* | 0.566 | 0.000 | 0.214 | 0.000 | -0.798 | 1 | -1.000 |
| Charge-Cationic* | -0.566 | 0.000 | -0.214 | 0.000 | 0.798 | -1.000 | 1 |
| Values in bold are different from 0 with a significance level of $\alpha = 0.05$ | | | | | | | |
| *Supplementary variables |  |  |  |  |  |  |  |

**Table 10.** Kaiser-Meyer-Olkin test for the sampling adequacy of factors and model for PCA.

| Variable | Viability | Concentration | MMP | Rigidity | ROS | Model |
| --- | --- | --- | --- | --- | --- | --- |
| KMO | 0.779 | 0.601 | 0.664 | 0.626 | 0.642 | 0.659 |

There was a significant effect of concentration, MMP, plasma membrane rigidity and ROS concentration on the viability of MDA MB 231 cells. ROS concentration had a significant negative correlation with the viability of the cells. However, there was no statistically significant correlation between ROS concentration and other factors. ROS concentration had a significant correlation with surface charge of nanoparticles with anionic AuNP having a negative correlation and cationic AuNP having a positive correlation.

Bartlett sphericity test was used to analyze the correlation matrix to test the null hypothesis that the correlation matrix is different from the identity matrix. It was observed that at least one of the correlations was significantly different than 0 with a p value of 0.0002. Further, Kaiser-Meyer-Olkin test (KMO test) was performed to measure the sampling adequacy of factors and the model for PCA analysis. KMO test is a statistical measure of the proportion of variance which is not attributed to common variance. Therefore, the data is suitable for PCA if KMO values are closer to 1. **Table** 10provides the KMO values for the factors and model. Based on KMO test, the viability of MDA MB 231 cell line was a middling factor, whereas, all other factors were mediocre factors. The overall model had a mediocre sampling adequacy. Therefore, we need more parameters to adequately define the effect of nanoparticle challenge on the viability of MDA MB 231 cells. However, we proceeded with the PCA analysis to understand the contribution of various parameters towards nanoparticle-mediated cytotoxic response.

The PCA was performed on the data, and six factors were identified (**Figure 13A**). The scree plot of percentage viability explained by each factor was used to identify two factors which could account for 91.43% of the variability of the data (**Figure 13B**). The correlation of various factors with principal components is given in **Figure 13C**, and the distribution of the data along two principal axes is provided in **Figure 13D**. The Factor 2 was able to distinguish between responses from anionic and cationic AuNP and had a strong correlation with the surface charge of the nanoparticle. The Factor 1 was able to differentiate between the concentration-dependent nanoparticle response and had a strong correlation with viability (Pearson’s correlation = 0.896), concentration of nanoparticles (Pearson’s correlation = -0.908), MMP (Pearson’s correlation = 0.917), and plasma membrane rigidity (Pearson’s correlation = 0.849). Therefore, all the factors used in PCA had a significant effect on the nanoparticle-mediated cytotoxic response.

**Figure 13.**
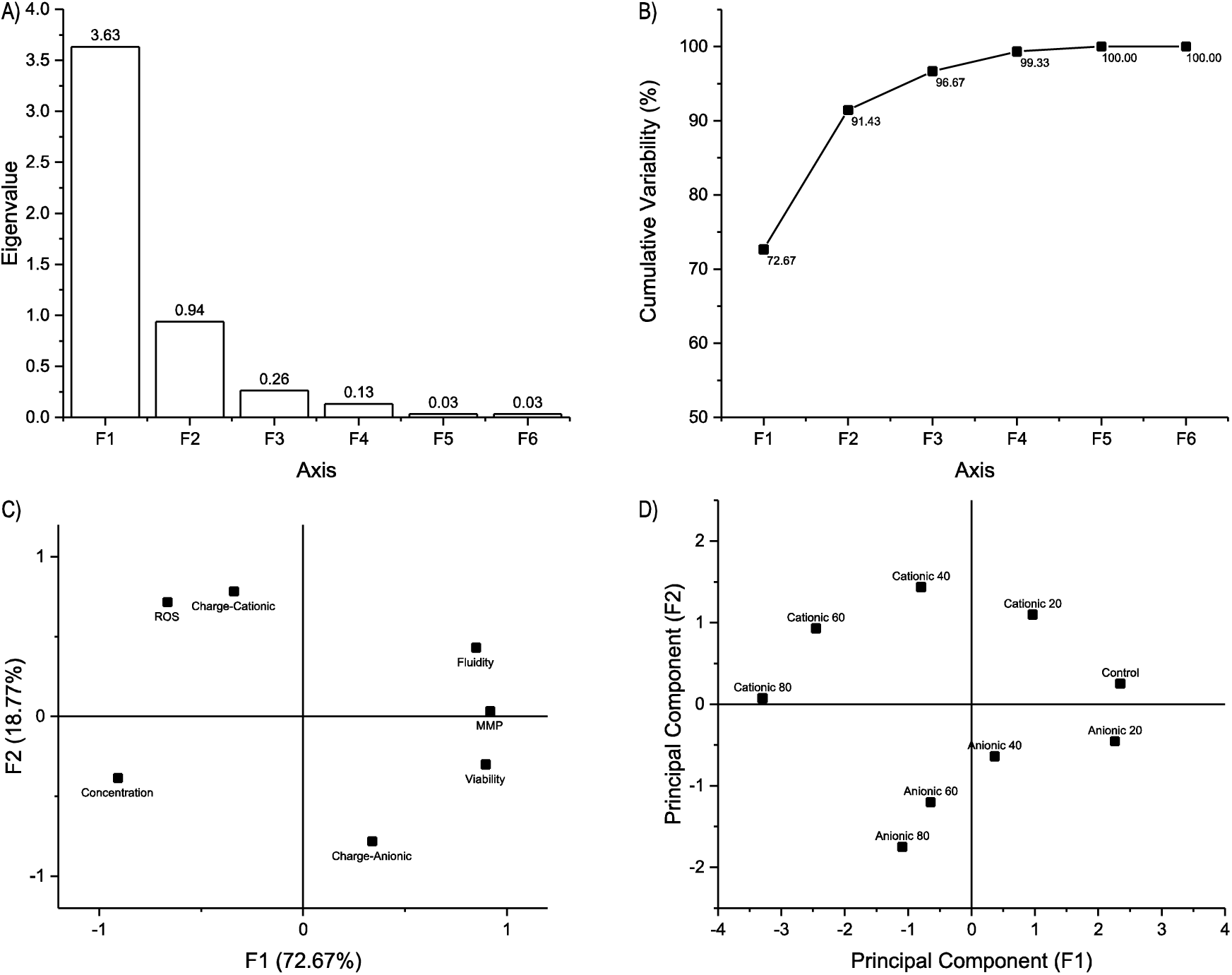
Principal Component Analysis (PCA) of cytotoxicity data. A) Scree plot of factors derived for the data, B) Cumulative variation explained by the factors; C) Correlation between variables and factors, and D) Scattered plot of data along the two principal factors.

### Model for cytotoxic response of AuNP

The data presented in this chapter was used to propose a generalized model for nanoparticle-mediated cytotoxic response. **Figure 14** provides the framework of the proposed model. The proposed model links the nanoparticle interaction mediated changes in the plasma membrane to the cellular fate. Nanoparticles adsorb biomolecules from the culture media, and the resulting biomolecular corona provides a unique identity to nanoparticles. The anionic and cationic AuNP tend to interact with the cell surface in a charge dependent manner as shown in section 0. Cationic AuNP cause a significant change in the cell surface charge at higher concentrations. The major change, however, was observed in the plasma membrane fluidity of the cells (section 0). The cationic nanoparticle caused a concentration-dependent increase in the lateral diffusivity of the plasma membrane. However, there is no report linking nanoparticle-mediated plasma membrane changes to the cellular fate.

**Figure 14.**
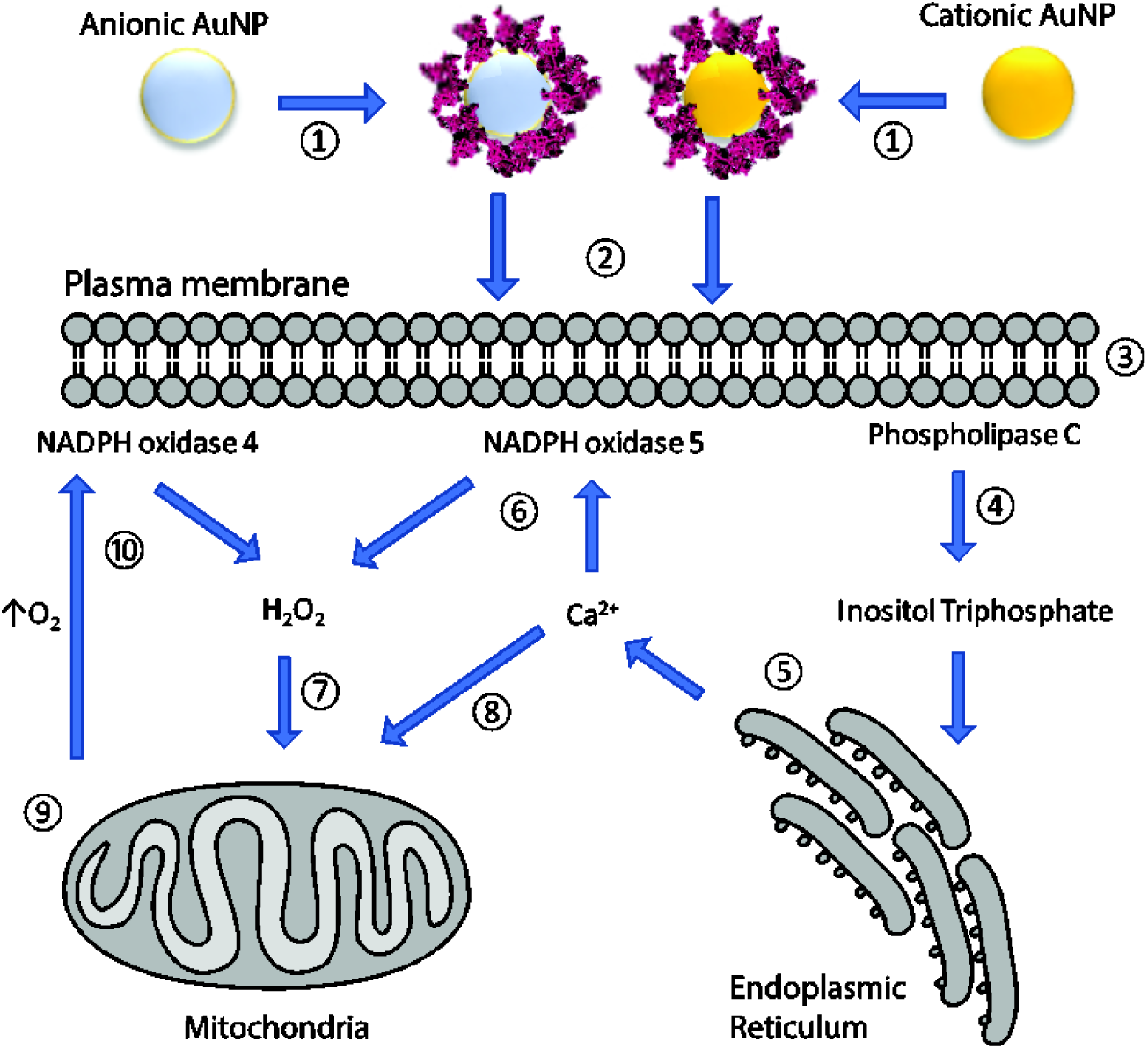
Proposed model for nanoparticle-mediated cytotoxic response. ① Nanoparticles adsorb proteins and other biomacromolecules from the biological milieu and form a corona around nanoparticles. ② Biomolecular corona provides a unique identity to the nanoparticle and is instrumental in establishing an interface between cells and nanoparticles. ③ Nanoparticles interact with the plasma membrane and alter the physicochemical properties of plasma membrane especially plasma membrane fluidity. ④ Alteration in plasma membrane fluidity activates phospholipase C which acts on phosphatidylinositol 4,5-bisphosphate (PIP2) and cleaves it into diacylglycerol (DAG) and inositol 1,4,5-trisphosphate (IP3). ⑤ IP3 is a soluble molecule and thus diffuses readily through the cytoplasm. On reaching ER, IP3 binds to the IP3 receptor which gates Ca^2+^ channels causing a release of Ca^2+^ from ER. ⑥ Ca^2+^ activates NADPH oxidase 5 (NOX5) resulting in the formation of superoxide ions. Intracellular superoxide dismutase converts superoxide to hydrogen peroxide. NOX are membrane bound enzymes and are present in both plasma membrane and ER. ⑦ NOX-mediated ROS cause activation of ATP-dependent potassium channels on mitochondrial membrane causing loss of mitochondrial membrane potential. Depolarized mitochondrial membrane causes the release of mitochondrial ROS to the cytoplasm resulting in an elevated oxidative stress. ⑧ Mitochondria also acts as an intracellular buffer for cytosolic Ca^2+^. Cytosolic Ca^2+^ enter the mitochondrial matrix through voltage-dependent calcium channels and calcium uniporters. ⑨ the mitochondrial overloading of Ca^2+^ elevates the respiratory status of mitochondria causing an increase in the cytosolic oxidative stress. The mitochondrial membrane also gets compromised as a result of opening of mitochondrial permeability transition pores (mPTP) causing a release of proapoptotic molecules into the cytosol leading to apoptotic cell death. ⑩ Loss of mitochondrial respiration due to mPTP opening also results in increased cytosolic oxygen level which is known to activate NOX4 resulting in elevated oxidative stress in the cell. In the case of excessive oxidative stress, cells can undergo necrosis.

Physicochemical characteristics of plasma membrane play a significant role in environmental stress sensing and signal transduction. The plasma membrane fluidity plays a vital role in cellular responses to heat stress^16^. Hyperfluidization of the plasma membrane during heat shock response in cells causes an increase in the cytosolic Ca^2+^ concentration both in calcium-free and calcium-containing media leading to mitochondrial hyperpolarization^53^. Therefore, the source of Ca^2+^ ions is intracellular and the major intracellular Ca^2+^ store is endoplasmic reticulum (ER). Heat shock also increases the phospholipase C activity^41^. Phospholipase C cleaves phosphatidylinositol-4,5-bisphosphate (PIP2) into diacylglycerol (DAG) and inositol-1,4,5-triphosphate (IP3). IP3 is a small molecule, and it diffuses freely into the cytoplasm. IP3 binds to the IP3 receptor (IP3R) present on the ER membrane. IP3R gates a Ca^2+^ channel and on binding with IP3, it releases the Ca^2+^ from the ER^54^. ***Therefore, we hypothesize that the nanoparticle-mediated changes in the plasma membrane fluidity would mimic heat stress response by plasma membrane causing an increase in the intracellular Ca***^***2+***^. A nanoparticle charge dependent transient increase in the intracellular Ca^2+^ has been reported for cationic AuNP challenge, whereas, neutral and anionic AuNP did not show any such effect^7^.

Cytosolic Ca^2+^ is a secondary messenger, and spatiotemporal changes in the concentration of Ca2^+^ are known to regulate cellular pathways involved in cell motility, proliferation, differentiation, endocytosis, exocytosis, and death (apoptosis and necrosis) to name a few^55^. Ca^2+^ ions also activate NADPH oxidase 5 (NOX5) which is a membrane-bond enzyme, and is localized primarily on the plasma membrane and ER membrane^56^. NOX 5 releases superoxide ions into the cytosol where superoxide dismutase converts superoxide ions into hydrogen peroxide (H_2_O_2_). H_2_O_2_ is one of the major intracellular ROS which increases the overall oxidative stress in the cell.

Mitochondria also act as an intracellular buffer for maintaining the cytosolic Ca^2+^ concentration^57^. Mitochondria actively take up Ca^2+^ by voltage sensitive calcium channels present on the outer mitochondrial membrane and ruthenium red-sensitive calcium uniporters situated in the inner mitochondrial membrane. Mitochondrial Ca^2+^ has a multitude of effect on cell physiology^58^. Mitochondria show an increase in activity as a result of Ca^2+^ uptake with an increase in the cytosolic concentration of NADPH and ATP. Transient opening of mitochondrial permeability transition pores (mPTP) maintains the mitochondrial Ca^2+^ levels. However, a prolonged opening of mPTP results in adverse effects on cellular physiology. A sustained increase in mitochondrial Ca^2+^ results in the loss of MMP leading to ATP depletion causing autophagic and necrotic responses, whereas, prolonged mPTP opening cause an increase in pro-apoptotic signals in the cytosol causing apoptotic or necrotic cell death^59^.

The increase in the cytosolic concentration of H_2_O_2_ also results in the decrease in MMP as a consequence of the activation of ATP-sensitive potassium channels present in the mitochondrial inner membrane^60^. The decrease in MMP results in loss of proton motive force (PMF) which is essential for the generation of ATP by the oxidative phosphorylation taking place on the mitochondrial membrane. This PMF uncoupling results in loss of mitochondrial oxygen uptake. NADPH oxidase 4 (NOX4) acts as an intracellular oxygen sensor, and NOX4 on activation produces H_2_O_2_^61^. Therefore, ***we propose that a mechanism based on NOX-mediated ROS generation along with the loss of mitochondrial bioactivity and sustained mPTP opening could explain the nanoparticle-mediated cytotoxic response and that this response stems from the physicochemical changes of plasma membrane caused by nanoparticle-cell interaction***.

MDA MB 231 cells used in this study have a high NOX activity^62^. NOX4 is relatively more abundant as compared to NOX5, whereas, the expression of NOX4 is relatively lower in SKBR3 as compared to MDA MB 231 cell line^63,64^. Therefore, NOX could potentially play a role in the observed cytotoxic response after nanoparticle challenge. The proposed model provides a link between nanoparticle-cell interaction and oxidative stress of cells resulting in a cytotoxic response which is mediated by the increase in intracellular calcium and loss of mitochondrial bioactivity. Cells respond to these stress conditions by undergoing proliferation at mild oxidative stress, apoptosis at intermediate oxidative stress and necrosis at high oxidative stress. Therefore, an increase in the intracellular Ca^2+^ could be linked with the cytotoxic response of nanoparticles. As evidence to the proposed model, cationic AuNP causes a significant increase in the intracellular Ca^2+^ and have a cytotoxic effect on cells^7^. In addition to cytotoxic response, the intracellular Ca^2+^ also plays a role in the differentiation of mesenchymal stem cells and migration of cancer cells. Therefore, ***the proposed model which establishes a link between nanoparticle-cell interaction and its role in modulating the oxidative stress by intracellular Ca***^***2+***^ ***can explain the plurality of nanoparticle-mediated responses in different cells.***

## Summary and conclusions

To summarize, two breast cancer cell lines (MDA MB 231 and SKBR3) were challenged with anionic and cationic AuNP, and it was observed that only cationic AuNP cause cytotoxicity to both cell lines in a concentration-dependent manner. Moreover, there was no significant difference between the response of cell lines to the AuNP challenge. Further, we observed that AuNP interaction with the plasma membrane of MDA MB 231 cells showed a surface charge dependent response. Cationic AuNP caused a significant change in the integrity, fluidity and surface charge of the plasma membrane, whereas, anionic AuNP had no such effect. Moreover, we also observed that cationic AuNP challenge resulted in significantly higher levels of cytosolic ROS as compared to anionic AuNP. However, incubation of both nanoparticles caused similar changes in the MMP indicating that mitochondria were not the source of ROS. Finally, PCA was performed to understand the influence of different parameters on cellular viability. There was a strong correlation between cell viability and other parameters including the concentration of nanoparticles, MMP, plasma membrane rigidity, and cytosolic ROS levels. Therefore, a generalized nanocytotoxicity model was proposed to include contributions from all parameters.

To conclude, the proposed model links physical interaction between nanoparticles and cells to a biological response of the cell. The modulation of intracellular levels of Ca^2+^ as a result of changes in the plasma membrane fluidity forms the basis of the proposed model. This model can also explain other biological effects of cells as a result of nanoparticle challenge as Ca^2+^ plays a significant role in a multitude of cellular processes including cell proliferation, differentiation, death, and signal transduction to name a few. Therefore, this model once proved, can help explain the observed plurality of cellular response to the nanoparticle challenge.

## References

1 Albanese, A., Tang, P. S. and Chan, W. C. (2012) The effect of nanoparticle size, shape, and surface chemistry on biological systems. Annual Review of Biomedical Engineering 14, 1–16.

2 Cedervall, T., Lynch, I., Lindman, S., Berggård, T., Thulin, E., Nilsson, H., Dawson, K. A. and Linse, S. (2007) Understanding the nanoparticle-protein corona using methods to quntify exchange rates and affinities of proteins for nanoparticles. Proceedings of the National Academy of Sciences of the United States of America 104, 2050-2055.

3 Lundqvist, M., Stigler, J., Elia, G., Lynch, I., Cedervall, T. and Dawson, K. A. (2008) Nanoparticle size and surface properties determine the protein corona with possible implications for biological impacts. Proceedings of the National Academy of Sciences of the United States of America 105, 14265-14270.

4 Raesch, S. S., Tenzer, S., Storck, W., Rurainski, A., Selzer, D., Ruge, C. A., Perez-Gil, J., Schaefer, U. F. and Lehr, C.-M. (2015) Proteomic and lipidomic analysis of nanoparticle corona upon contact with lung surfactant reveals differences in protein, but not lipid composition. ACS Nano 9, 11872-11885.

5 Wan, S., Kelly, P. M., Mahon, E., Stöckmann, H., Rudd, P. M., Caruso, F., Dawson, K. A., Yan, Y. and Monopoli, M. P. (2015) The “sweet” side of the protein corona: Effects of glycosylation on nanoparticle–cell interactions. ACS Nano 9, 2157-2166.

6 Walczyk, D., Bombelli, F. B., Monopoli, M. P., Lynch, I. and Dawson, K. A. (2010) What the Cell “Sees” in Bionanoscience. Journal of the American Chemical Society 132, 5761-5768.

7 Arvizo, R. R., Miranda, O. R., Thompson, M. A., Pabelick, C. M., Bhattacharya, R., Robertson, J. D., Rotello, V. M., Prakash, Y. S. and Mukherjee, P. (2010) Effect of Nanoparticle Surface Charge at the Plasma Membrane and Beyond. Nano Letters 10, 2543-2548.

8 Yi, C., Liu, D., Fong, C.-C., Zhang, J. and Yang, M. (2010) Gold Nanoparticles Promote Osteogenic Differentiation of Mesenchymal Stem Cells through p38 MAPK Pathway. ACS Nano 4, 6439-6448.

9 Fatisson, J., Quevedo, I. R., Wilkinson, K. J. and Tufenkji, N. (2012) Physicochemical characterization of engineered nanoparticles under physiological conditions: Effect of culture media components and particle surface coating. Colloids and Surfaces B: Biointerfaces 91, 198-204.

10 Huang, D., Zhou, H., Liu, H. and Gao, J. (2015) The cytotoxicity of gold nanoparticles is dispersity-dependent. Dalton Transactions 44, 17911-17915.

11 Huang, D., Zhou, H. and Gao, J. (2015) Nanoparticles modulate autophagic effect in a dispersity-dependent manner. Scientific Reports 5, 14361

12 Nel, A. E., Mädler, L., Velegol, D., Xia, T., Hoek, E. M. V., Somasundaran, P., Klaessig, F., Castranova, V. and Thompson, M. (2009) Understanding biophysicochemical interactions at the nano-bio interface. Nature Materials 8, 543-557.

13 Peetla, C., Jin, S., Weimer, J., Elegbede, A. and Labhasetwar, V. (2014) Biomechanics and thermodynamics of nanoparticle interactions with plasma and endosomal membrane lipids in cellular uptake and endosomal escape. Langmuir 30, 7522-7532.

14 Wang, S.-H., Lee, C.-W., Chiou, A. and Wei, P.-K. (2010) Size-dependent endocytosis of gold nanoparticles studied by three-dimensional mapping of plasmonic scattering images. Journal of Nanobiotechnology 8, 33.

15 Zhang, Y., Yang, M., Portney, N. G., Cui, D., Budak, G., Ozbay, E., Ozkan, M. and Ozkan, C. S. (2008) Zeta potential: A surface electrical characteristic to probe the interaction of nanoparticles with normal and cancer human breast epithelial cells. Biomedical Microdevices 10, 321-328.

16 Balogh, G., Horváth, I., Nagy, E., Hoyk, Z., Benkõ, S., Bensaude, O. and Vígh, L. (2005) The hyperfluidization of mammalian cell membranes acts as a signal to initiate the heat shock protein response. FEBS Journal 272, 6077-6086.

17 Nagy, E., Balogi, Z., Gombos, I., Åkerfelt, M., Björkbom, A., Balogh, G., Török, Z., Maslyanko, A., Fiszer-Kierzkowska, A. and Lisowska, K. (2007) Hyperfluidization-coupled membrane microdomain reorganization is linked to activation of the heat shock response in a murine melanoma cell line. Proceedings of the National Academy of Sciences 104, 7945-7950.

18 Török, Z., Crul, T., Maresca, B., Schütz, G. J., Viana, F., Dindia, L., Piotto, S., Brameshuber, M., Balogh, G., Péter, M., Porta, A., Trapani, A., Gombos, I., Glatz, A., Gungor, B., Peksel, B., Vigh Jr, L., Csoboz, B., Horváth, I., Vijayan, M. M., Hooper, P. L., Harwood, J. L. and Vigh, L. (2014) Plasma membranes as heat stress sensors: From lipid-controlled molecular switches to therapeutic applications. Biochimica et Biophysica Acta (BBA) - Biomembranes 1838, 1594-1618.

19 Zhou, Y., Wong, C.-O., Cho, K.-j., van der Hoeven, D., Liang, H., Thakur, D. P., Luo, J., Babic, M., Zinsmaier, K. E. and Zhu, M. X. (2015) Membrane potential modulates plasma membrane phospholipid dynamics and K-Ras signaling. Science 349, 873-876.

20 Page, B., Page, M. and Noel, C. (1993) A new fluorometric assay for cytotoxicity measurements in vitro. International Journal of Oncology 3, 473-473.

21 Jones, K. H. and Senft, J. A. (1985) An improved method to determine cell viability by simultaneous staining with fluorescein diacetate-propidium iodide. Journal of Histochemistry & Cytochemistry 33, 77-79.

22 Galla, H. J. and Sackmann, E. (1974) Lateral diffusion in the hydrophobic region of membranes: use of pyrene excimers as optical probes. Biochimica et Biophysica Acta 339, 103-115.

23 Scaduto, R. C. and Grotyohann, L. W. (1999) Measurement of mitochondrial membrane potential using fluorescent rhodamine derivatives. Biophysical Journal 76, 469-477.

24 Wang, H. and Joseph, J. A. (1999) Quantifying cellular oxidative stress by dichlorofluorescein assay using microplate reader. Free Radical Biology and Medicine 27, 612-616.

25 Hinderliter, P. M., Minard, K. R., Orr, G., Chrisler, W. B., Thrall, B. D., Pounds, J. G. and Teeguarden, J. G. (2010) ISDD: A computational model of particle sedimentation, diffusion and target cell dosimetry for in vitro toxicity studies. Particle and Fibre Toxicology 7, 36.

26 Cho, E. C., Zhang, Q. and Xia, Y. (2011) The effect of sedimentation and diffusion on cellular uptake of gold nanoparticles. Nature Nanotechnology 6, 385-391.

27 Ahmad Khanbeigi, R., Kumar, A., Sadouki, F., Lorenz, C., Forbes, B., Dailey, L. A. and Collins, H. (2012) The delivered dose: Applying particokinetics to in vitro investigations of nanoparticle internalization by macrophages. Journal of Controlled Release 162, 259-266.

28 Zook, J. M., MacCuspie, R. I., Locascio, L. E., Halter, M. D. and Elliott, J. T. (2011) Stable nanoparticle aggregates/agglomerates of different sizes and the effect of their size on hemolytic cytotoxicity. Nanotoxicology 5, 517-530.

29 Halamoda-Kenzaoui, B., Ceridono, M., Colpo, P., Valsesia, A., Urbán, P., Ojea-Jiménez, I., Gioria, S., Gilliland, D., Rossi, F. and Kinsner-Ovaskainen, A. (2015) Dispersion behaviour of silica nanoparticles in biological media and its influence on cellular uptake. PloS one 10, e0141593..

30 Reddy, A., Caler, E. V. and Andrews, N. W. (2001) Plasma Membrane Repair Is Mediated by Ca2+-Regulated Exocytosis of Lysosomes. Cell 106, 157-169.

31 Huynh, C., Roth, D., Ward, D. M., Kaplan, J. and Andrews, N. W. (2004) Defective lysosomal exocytosis and plasma membrane repair in Chediak–Higashi/beige cells. Proceedings of the National Academy of Sciences of the United States of America 101, 16795-16800.

32 Oh, N. and Park, J. H. (2014) Surface Chemistry of Gold Nanoparticles Mediates Their Exocytosis in Macrophages. Acs Nano 8, 6232-6241.

33 Ruenraroengsak, P., Novak, P., Berhanu, D., Thorley, A. J., Valsami-Jones, E., Gorelik, J., Korchev, Y. E. and Tetley, T. D. (2012) Respiratory epithelial cytotoxicity and membrane damage (holes) caused by amine-modified nanoparticles. Nanotoxicology 6, 94-108.

34 Goldenberg, N. M. and Steinberg, B. E. (2010) Surface charge: a key determinant of protein localization and function. Cancer Research 70, 1277-1280.

35 Su, G., Zhou, H., Mu, Q., Zhang, Y., Li, L., Jiao, P., Jiang, G. and Yan, B. (2012) Effective Surface Charge Density Determines the Electrostatic Attraction between Nanoparticles and Cells. The Journal of Physical Chemistry C 116, 4993-4998.

36 Balogh, G., Maulucci, G., Gombos, I., Horváth, I., Török, Z., Péter, M., Fodor, E., Páli, T., Benkő, S. and Parasassi, T. (2011) Heat stress causes spatially-distinct membrane re-modelling in K562 leukemia cells. PloS one 6, e21182..

37 Balogh, G. Role of membranes in mammalian stress response: sensing, lipid signals and adaptation, Cardiff University, (2011).

38 Chabanel, A., Abbott, R. E., Chien, S. and Schachter, D. (1985) Effects of benzyl alcohol on erythrocyte shape, membrane hemileaflet fluidity and membrane viscoelasticity. Biochimica et Biophysica Acta (BBA)-Biomembranes 816, 142-152.

39 Wang, B., Zhang, L., Sung, C. B. and Granick, S. (2008) Nanoparticle-induced surface reconstruction of phospholipid membranes. Proceedings of the National Academy of Sciences of the United States of America 105, 18171-18175.

40 Pompa, P. P., Vecchio, G., Galeone, A., Brunetti, V., Sabella, S., Maiorano, G., Falqui, A., Bertoni, G. and Cingolani, R. (2011) In vivo toxicity assessment of gold nanoparticles in Drosophila melanogaster. Nano Research 4, 405-413.

41 Calderwood, S. K. and Stevenson, M. A. (1993) Inducers of the heat shock response stimulate phospholipase C and phospholipase A2 activity in mammalian cells. Journal of cellular physiology 155, 248-256.

42 Hsu, Y.-L., Yu, H.-S., Lin, H.-C., Wu, K.-Y., Yang, R.-C. and Kuo, P.-L. (2011) Heat shock induces apoptosis through reactive oxygen species involving mitochondrial and death receptor pathways in corneal cells. Experimental Eye Research 93, 405-412.

43 Nel, A., Xia, T., Mädler, L. and Li, N. (2006) Toxic potential of materials at the nanolevel. Science 311, 622-627.

44 Li, N., Xia, T. and Nel, A. E. (2008) The role of oxidative stress in ambient particulate matter-induced lung diseases and its implications in the toxicity of engineered nanoparticles. Free Radical Biology and Medicine 44, 1689-1699.

45 Aw, T. Y. (1999) Molecular and cellular responses to oxidative stress and changes in oxidation-reduction imbalance in the intestine. The American Journal of Clinical Nutrition 70, 557-565.

46 Turrens, J. F. (2003) Mitochondrial formation of reactive oxygen species. The Journal of Physiology 552, 335-344.

47 Moon, E. J., Sonveaux, P., Porporato, P. E., Danhier, P., Gallez, B., Batinic-Haberle, I., Nien, Y.-C., Schroeder, T. and Dewhirst, M. W. (2010) NADPH oxidase-mediated reactive oxygen species production activates hypoxia-inducible factor-1 (HIF-1) via the ERK pathway after hyperthermia treatment. Proceedings of the National Academy of Sciences of the United States of America 107, 20477-20482.

48 Schulz, E., Wenzel, P., Münzel, T. and Daiber, A. (2014) Mitochondrial redox signaling: interaction of mitochondrial reactive oxygen species with other sources of oxidative stress. Antioxidants & redox signaling 20, 308-324.

49 Mateo, D., Morales, P., Ávalos, A. and Haza, A. I. (2014) Oxidative stress contributes to gold nanoparticle-induced cytotoxicity in human tumor cells. Toxicology Mechanisms and Methods 24, 161-172.

50 Arvizo, R. R., Saha, S., Wang, E., Robertson, J. D., Bhattacharya, R. and Mukherjee, P. (2013) Inhibition of tumor growth and metastasis by a self-therapeutic nanoparticle. Proceedings of the National Academy of Sciences 110, 6700-6705.

51 Zhang, D., Liu, D., Zhang, J., Fong, C. and Yang, M. (2014) Gold nanoparticles stimulate differentiation and mineralization of primary osteoblasts through the ERK/MAPK signaling pathway. Materials Science and Engineering: C 42, 70-77.

52 Huang, H., Quan, Y.-y., Wang, X.-p. and Chen, T.-s. (2016) Gold nanoparticles of diameter 13 nm induce apoptosis in rabbit articular chondrocytes. Nanoscale research letters 11, 249.

53 Rusnak, F. and Mertz, P. (2000) Calcineurin: form and function. Physiological Reviews 80, 1483-1521.

54 Mikoshiba, K. (2007) IP3 receptor/Ca^2+^ channel: from discovery to new signaling concepts. Journal of neurochemistry 102, 1426-1446.

55 Monteith, G. R., McAndrew, D., Faddy, H. M. and Roberts-Thomson, S. J. (2007) Calcium and cancer: targeting Ca2+ transport. Nature Reviews Cancer 7, 519-530.

56 Brown, D. I. and Griendling, K. K. (2009) Nox proteins in signal transduction. Free Radical Biology and Medicine 47, 1239-1253.

57 Rizzuto, R., De Stefani, D., Raffaello, A. and Mammucari, C. (2012) Mitochondria as sensors and regulators of calcium signalling. Nature Reviews Molecular Cell Biology 13, 566-578.

58 Rasola, A. and Bernardi, P. (2011) Mitochondrial permeability transition in Ca^2+^-dependent apoptosis and necrosis. Cell calcium 50, 222-233.

59 Kinnally, K. W., Peixoto, P. M., Ryu, S.-Y. and Dejean, L. M. (2011) Is mPTP the gatekeeper for necrosis, apoptosis, or both? Biochimica et Biophysica Acta (BBA)-Molecular Cell Research 1813, 616-622.

60 Facundo, H. T., de Paula, J. G. and Kowaltowski, A. J. (2007) Mitochondrial ATP-sensitive K^+^ channels are redox-sensitive pathways that control reactive oxygen species production. Free Radical Biology and Medicine 42, 1039-1048.

61 Nisimoto, Y., Diebold, B. A., Cosentino-Gomes, D. and Lambeth, J. D. (2014) Nox4: a hydrogen peroxide-generating oxygen sensor. Biochemistry 53, 5111-5120.

62 Choi, J., Jung, Y., Kim, J., Kim, H. and Lim, I. (2016) Inhibition of breast cancer invasion by TIS21/BTG2/Pc3-Akt1-Sp1-Nox4 pathway targeting actin nucleators, mDia genes. Oncogene 35, 83-93.

63 Choi, J.-A., Lee, J.-W., Kim, H., Kim, E.-Y., Seo, J.-M., Ko, J. and Kim, J.-H. (2010) Pro-survival of estrogen receptor-negative breast cancer cells is regulated by a BLT2–reactive oxygen species-linked signaling pathway. Carcinogenesis 31, 543-551.

64 Graham, K. A., Kulawiec, M., Owens, K. M., Li, X., Desouki, M. M., Chandra, D. and Singh, K. K. (2010) NADPH oxidase 4 is an oncoprotein localized to mitochondria. Cancer Biology & Therapy 10, 223-231.

